# Unique features of transcription termination and initiation at closely spaced tandem human genes

**DOI:** 10.1101/2021.09.10.459726

**Authors:** Noa Nissani, Igor Ulitsky

## Abstract

The synthesis of RNA Polymerase II (Pol2) products, which include messenger RNAs or long noncoding RNAs, culminates in transcription termination. How the transcriptional termination of a gene impacts the activity of promoters found immediately downstream of it, and which can be subject to potential transcriptional interference, remains largely unknown. We examined in an unbiased manner features of the intergenic region of pairs of tandem and closely spaced (<2kb) genes found on the same strand. Intergenic regions separating tandem genes are enriched with Guanines and are characterized by binding of several proteins, including AGO1 and AGO2 of the RNA interference pathway. Additionally, we found that Pol2 with a specific modification pattern is particularly enriched in this region, and it is lost upon perturbations affecting splicing or transcriptional elongation. Perturbations of genes involved in Pol2 pausing and R loop biology preferentially affect expression of downstream genes in tandem gene pairs. Overall, we find that features associated with potential Pol2 recycling rather than those associated with avoidance of transcriptional interference are the predominant driving force shaping these regions.

## Introduction

Transcription of messenger RNAs (mRNAs) and long noncoding RNAs (lncRNAs) begins with the heavily regulated transcription initiation where RNA Polymerase 2 (Pol2) complex is assembled on promoters. It proceeds with transcriptional elongation, which is often coordinated with processing of the nascent RNA (capping its 5’ and splicing out the introns), and culminates with 3’ end formation, which almost universally involves cleavage and polyadenylation (CPA) of the nascent RNA transcript following a polyadenylation signal (PAS) and the addition of a poly-A tail. As opposed to transcription initiation, the mechanisms of transcription termination are relatively less understood (Gruber and Zavolan 2019).

It is important to distinguish between the site of CPA, which defines the end of the transcript and the site of Pol2 release from the DNA, since Pol2 dissociates from the DNA between 100bp to several kb downstream of the polyadenylation site (Ashfield, Enriquez-Harris, and Proudfoot 1991a; Dye and Proudfoot 2001; Hagenbüchle et al. 1984; Tantravahi, Alvira, and Falck-Pedersen 1993). Disengagement of Pol2 from the DNA is thought to be important for both Pol2 recycling into the cellular pool and for insulating elongating Pol2 from downstream promoters to allow proper initiation at the downstream gene (Greger and Proudfoot 1998; Gromak, West, and Proudfoot 2006).

There are two main models for the dissociation of Pol2 from the DNA: the allosteric model and the torpedo model. The allosteric model proposes that once the elongating Pol2 passes over a functional PAS, it undergoes a conformational change. This conformational change is thought to be mediated by the association of CPA factors with Pol2 CTD which results in a Pol2 pause and its eventual release. The torpedo model connects between nascent RNA cleavage step and Pol2 release. In this model, XRN2, as part of the 5’-to-3’ RNA degradation machinery, engages with the free 5’ formed on the nascent transcript that is still being synthesized by Pol2 following CPA. XRN2 digests this RNA faster than the speed of Pol2 elongation and thus acts as a torpedo until it reaches Pol2 and triggers its release from the DNA (Proudfoot 2016). Indeed, depletion of Xrn2 in mammalian cells or of its ortholog Rat1 in the yeast *Saccharomyces Cerevisiae* has a strong effect on transcriptional termination (Kim et al. 2004; West, Gromak, and Proudfoot 2004).

Having a paused, or slowed-down, Pol2 downstream of termination sites is not a feature uniquely ascribed to the allosteric model, but it may also enhance XRN2-mediated termination and promote Pol2 recycling. Paused Pol2 may be giving an advantage to XRN2 in this race. G-rich elements are thought to enhance Pol2 pausing, possibly through promoting the formation of R-loop structures (DNA:RNA hybrids) (Skourti-Stathaki, Proudfoot, and Gromak 2011). Potentially related, a G-rich sequence motif, the MAZ element (G5AG5), bound by MAZ transcription factor (TF), was also shown to be required for efficient termination between closely spaced human complement genes C2 and factor B, which are separated by a mere 421 bp (Ashfield, Enriquez-Harris, and Proudfoot 1991b; Ashfield et al. 1994). This sequence element was also shown to be sufficient for polyadenylation *in vitro*. In contrast, similarly G-rich Sp1 binding sites G5CG5 were not effective in this system. Pausing at such sequences was shown to be an intrinsic property of Pol2, at least *in vitro* (Yonaha and Proudfoot 1999).

The different transcription steps are orchestrated by modification of heptapeptide repeats of the free CTD of Pol2 largest subunit, RPB1 (Harlen and Churchman 2017). At the promoter, Pol2 is mostly unphosphorylated. During initial elongation, Serine-5 and Tyrosine-1 gradually become phosphorylated (S5P and Y1P), which might aid with recruitment of the RNA capping machinery. Later elongation stages are correlated with a decrease in S5P and an increase in S2P. Furthermore, S2P is associated with 3’ ends of genes and interacts with components of the CPA machinery. In addition, phosphorylated Threonine-4 (T4P) has also been correlated with termination regions (McCracken et al. 1997; Heidemann et al. 2013; Hsin and Manley 2012; Schlackow et al. 2017; Nojima et al. 2018).

While features of closely positioned promoters have been studied systematically (Chen et al. 2016), features of closely spaced termination and initiation regions have been less explored. The physical architecture and order of the mammalian genome consists of two predominant types of regions – gene-rich and gene-poor, and those are associated with overall differences in sequence composition and other features, such as intron length (Amit et al. 2012). Gene-rich regions are usually characterized by more euchromatic regions, which are thought to facilitate wide expression. A subset of these clustered genes are found in particularly close proximity to each other (<2kb distance). A further subset of these precede each other on the same strand, we refer to such gene pairs as “tandem genes”. The precision of the termination process is thought to have significant importance especially in the case of tandem genes, as it ensures the production of two stable transcripts, avoiding a readthrough transcript between a non-properly terminated upstream gene and continuing at the promoter of the downstream gene (Proudfoot 2016; Shearwin, Callen, and Egan 2005). In cases where both genes are well-expressed in the same cells, we expect that efficiency of termination is particularly important to allow both transcripts to be expressed at the right levels, and we hypothesize that this efficiency is encoded in the intergenic sequence. Notably, this intergenic sequence may optimize both avoidance of interference between the two transcripts, as well as potential recycling of the terminating Pol2 to be re-used in transcription of the downstream gene.

We became interested in the potential crosstalk between adjacent transcriptional units following our observation that in mouse embryonic fibroblasts, loss of the *Chaserr* lncRNA and the consequent increased dosage of CHD2 chromatin remodeler leads to repression of promoters found within <2kb of highly transcribed genes on the same strand (Rom et al. 2019). This observation suggested that there is potential for substantial transcriptional crosstalk between closely-spaced tandem genes in mammalian cells, echoing studies in yeast (Thebault et al. 2011; Martens, Laprade, and Winston 2004; Hainer et al. 2011; Pruneski et al. 2011) and in other species (Shuman 2020), and motivated us to look more broadly at features and perturbation sensitivities shared by such transcriptional units, which we describe below.

## Results

### Prevalence of closely spaced and co-expressed tandem gene pairs in the human genome

We first analyzed the overall prevalence of adjacent closely-spaced genes found on the same strand. We grouped pairs of adjacent human protein-coding genes, considering genes >5kb and <800kb in length, and removing all pairs of overlapping genes (see Methods), we then split them into groups based on their genomic orientation relative to each other: divergent genes, situated on different strands with transcription facing in opposite directions (2,737 pairs, 23.2%); convergent, pairs on different strands, with transcription directed towards each other (2,866 pairs, 24.3%); or tandem genes, transcribed from the same strand (6,177 pairs, 52.4%). The distribution of the relative orientations of genes in the human genome is thus roughly as expected by chance. Adjacent gene pairs were further separated into groups based on their distance from one another (defined as the minimal distance between transcribed bases). Interestingly, the proportional share of closely spaced genes (<2kb and 2–5kb) was generally higher than expected, with the exception of divergent genes separated by 2–5 kb, which were relatively depleted, whereas divergent genes in the <2kb category were relatively more common, as expected for genes that can share a common promoter (**Fig. 1A** and **Table S1**). Notably, 222 (49%) of the close tandem pairs, 285 (51%) of the close divergent pairs and 315 (60%) of the close convergent pairs have homologs that meet the same criteria in the mouse genome (see Methods), suggesting that close tandem spacing is not strongly avoided. We then focused on the set of tandem genes, and analyzed in detail 457 pairs of adjacent, closely spaced (<2kb) genes (with the constraints imposed above).. We further classified tandem pairs as co-expressed (188 pairs, ∼41%) based on HepG2 cell line expression data from the ENCODE project (Djebali et al. 2012) (we obtained very similar results when using ENCODE K562 data, and so focused on HepG2 in the rest of the analysis, unless indicated otherwise) (see Methods and **Tables S2-S9**). We analyzed the co-expression proportion in the other subgroups divided by orientations and distances and found that co-expression was most common in the closely spaced pairs of genes, regardless of their orientation (**Fig. 1A**). Furthermore, this trend remained if the distance between the genes was defined as the distance between their promoters (**Fig. 1B**). We conclude that compared to the convergent orientation, there are fewer tandem gene pairs separated by <2kb, perhaps because the A/T-rich polyadenylation signals are less likely to co-occur with G/C-rich promoters (see below). Nevertheless, when the distance between adjacent genes is short, there is no evidence that tandem genes are less likely to be co-expressed, which could be expected if transcriptional interference was posing a substantial obstruction to transcription from downstream promoters.

**Figure 1.**
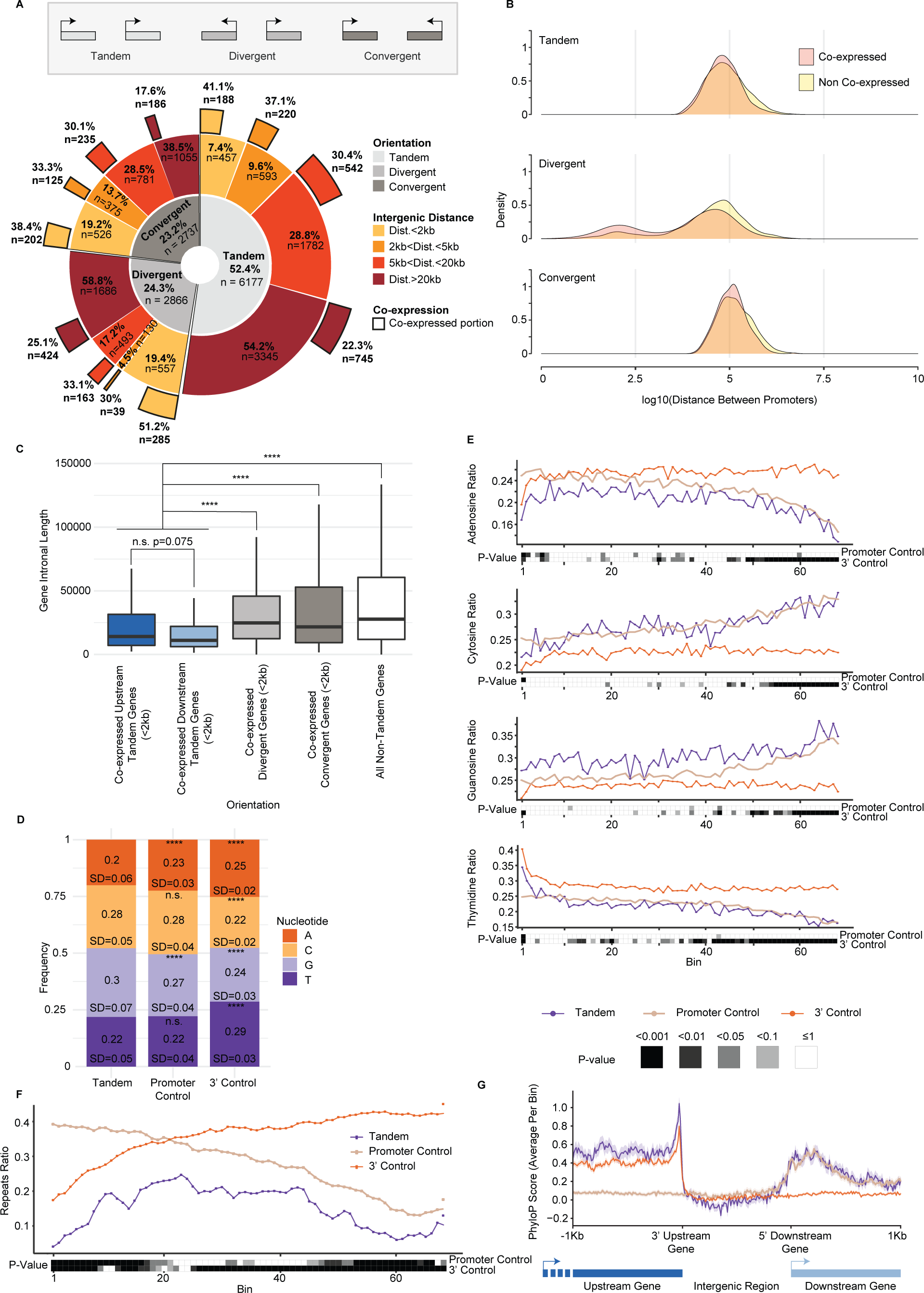
Genomic architecture and nucleotide composition of tandem genes. (A) Donut chart displaying the distribution of different gene orientations (inner circle, dataset of genes defined in the Methods section), distance between the genes (middle circle), and portion of co-expressed genes within each distance group (outer circle) defined in HepG2 cell line. (B) Density plot of the log10-transformed distance between the promoters of tandem (top), divergent (middle), or convergent (bottom) co-expressed (light red) or non-co-expressed (yellow) pairs of genes. Co-expression was tested in HepG2 cell line. (C) Boxplot of distribution of the sum of introns length for co-expressed genes in the various genomic orientations. Co-expression defined in HepG2. The thickened line represents the median intronal length of the genes, the lower and upper boxplot hinges correspond to first and third percentiles of the data, respectively. The whiskers represent the minimal/maximal existing value within 1.5 × inter-quartile range. Outliers were removed from the analysis. ****: P<=0.0001; Wilcoxon rank-sum test. (D) Overall nucleotide frequency within HepG2 co-expressed STIRs and within their promoter or 3’ controls. Shown are the standard deviation and paired Wilcoxon rank-sum test P-values between STIRs and either control. ****: P<=0.0001. (E) Metagene analysis showing binned nucleotide ratio within co-expressed STIRs (purple), promoter controls (beige) or 3’ controls (orange), of HepG2 cells. Heatmap shows the Bonferroni corrected P-value of paired Wilcoxon rank-sum test between STIRs and their respective controls within each bin. (F) Metagene analysis of transposable elements occupancy within HepG2 co-expressed STIRs or their respective controls. Bottom heatmap as in E. (G) Average PhyloP conservation scores within HepG2 co-expressed STIRs and flanking 1kb at the ends of the upstream (3’ end) and downstream (TSS) tandem genes (blue and teal, respectively). 5’ of promoter control genes are aligned to the 5’ downstream gene (beige) and the TTS of 3’ control genes are aligned to the 3’ upstream gene (orange).

### Sequence features of intergenic regions separating tandem co-expressed gene pairs

We reasoned that co-expressed pairs of closely spaced tandem genes, that need to be produced at similar conditions in many tissues, would require, at least in some tissues where they are highly expressed, a more precise regulation over the termination process. Additionally, we considered the possibility that the vicinity of the 3’ end of one gene to the promoter of another may facilitate the recycling of Pol2 machinery via some sequence or transcriptional features within the “short tandem intergenic region” (STIR). As a group, closely spaced co-expressed tandem genes tend to have substantially shorter introns than other genes (**Fig. 1C**), consistent with previous observations about differences between gene-rich and gene-poor regions of the human genome (Amit et al. 2012). Markedly, introns of co-expressed tandem genes are also shorter than those of co-expressed closely-spaced divergent or convergent genes. These differences dictated our strategy for selecting control genomic regions. As a control for the set of co-expressed adjacent tandem gene pairs, we matched for each pair of a gene A found upstream of B five “promoter-control” genes for B – these genes had a similar intronal length and expression levels as B (based on ENCODE expression data mentioned above), but no close upstream gene within at least 5kb. Similarly, we matched five “3’ control” genes for A – these genes had a similar intronal length and expression pattern to A, but no close downstream neighbor (**Fig. S1A** and Methods). Because the set of expressed genes differs between cell lines, the set of controls was also cell-line– specific. Importantly, in the following analyses, the lengths of the control regions we use are the same as those of the STIRs. For example, as controls for the 789 bp STIR between the genes *TULP1* and *TEAD3* we used 789 bp upstream of the promoters of five genes expressed similarly and with similar intronal length as TEAD3, and 789 bp regions downstream of the transcription termination sites (TTS) of five genes expressed similarly and with similar intronal length as TULP1.

STIRs were generally slightly more GC-rich than both types of control regions (58% G/C on average for STIRs compared to 46% and 55% at 3’ control regions and promoter regions in HepG2 cell line, respectively) (**Fig. 1D**). The nucleotide composition at STIRs was more similar to the promoter control regions than to the 3’ control regions (**Fig. 1E**). Notably, when compared to the promoter controls, there was an enrichment of Gs, spanning the full length of the STIR, and depletion of As in the beginning of the STIRs (**Fig. 1E**). Furthermore, STIRs were depleted of transposable elements compared to both controls (**Fig. 1F**). This may point to the functional importance of STIR sequences, but interestingly, when considering sequence conservation during vertebrate evolution, we observed slightly lower conservation levels of STIRs compared to control sequences. Notably, the 3’ region of the upstream co-expressed tandem gene shows some preferential conservation compared to its 3’ control gene set (**Fig. 1G**). The proximity of a CPA site upstream of a promoter on the same strand thus seems to have a mild effect on the sequence composition in the STIR and on evolutionary conservation at the upstream CPA site.

### G-rich sequence motifs are enriched in STIRs

The observation that STIRs have a biased sequence composition prompted us to look for specific enriched motifs within these regions. To this end, we used STREME (Sensitive, Thorough, Rapid, Enriched Motif Elicitation) from the MEME suite package (Bailey et al. 2015; Bailey 2011, 2021) and filtered for motifs both enriched within STIRs compared to the background model (as defined by STREME) and more prevalent in STIRs relative to promoter and 3’ control regions using FIMO (Grant, Bailey, and Noble 2011) (see Methods). We performed the enrichment analysis using both an “RNA” mode that seeks motifs only in the strand where both genes are transcribed, and a “DNA” mode, in which motifs are sought on both strands. Our analysis highlighted GGGGCGGG and GGGGCGGGGSC found in RNA (STREME P=0.023 and P=0.035, respectively) and CCTTCCC found in DNA (P=0.02) modes (**Fig. 2A** **and S2A**) as the lead motifs found in ∼33%, ∼49% and ∼43% of the STIRs, as opposed to ∼22%, ∼33% and ∼30% of control promoter regions and ∼3%, ∼15% and 23% of the 3’ controls, and with ∼0.53, ∼0.97 and ∼0.43 average motif occurrences in STIRs as opposed to averages of ∼0.36, ∼0.74 and ∼0.3 or ∼0.04, ∼0.21 and ∼0.23 in the promoter and 3’ controls, respectively. Notably, most enriched motifs in this analysis are particularly G/C-rich (**Fig. 2B** and **S2B-C**). Moreover, examination of the GGGGNGGGG motifs across the STIRs of co-expressed pairs in HepG2 and K562 showed enrichment in STIRs over both types of controls. As expected, we found enrichment of the ‘GGGGCGGGG’ variant, which is very similar to the ‘GGGGCGGG’ motif identified *de novo* by STREME. ‘GGGGTGGGG’ and to some extent ‘GGGGAGGGG’ were also enriched in STIRs. Intriguingly, this was not the case for ‘GGGGGGGGG’ which was relatively depleted from all sequences inspected, and further depleted in STIRs.

**Figure 2.**
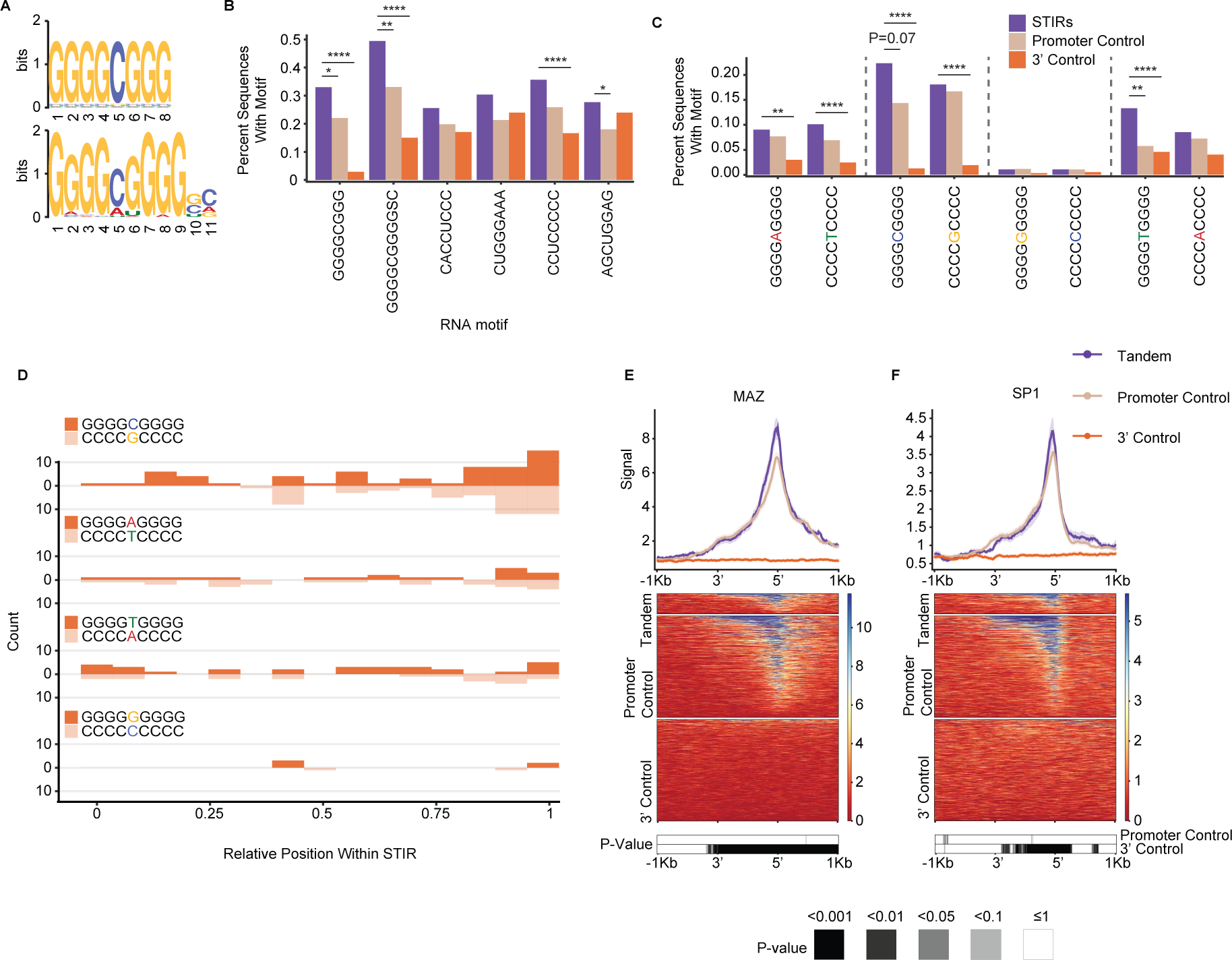
Enriched motifs in STIRs. (A) Logo representation of motifs identified by STREME in the “RNA” mode as enriched in either HepG2 (top) or K562 (bottom) co-expressed STIRs. (B) Barplots showing the proportion of STIRs (purple) or control sequences (beige and orange) that carry the RNA-mode STREME-discovered motif. Shown are Bonferroni corrected proportion test P-values (*:P<=0.05, **:P<=0.01, ***:P<=0.001, ****<=0.0001). (C) Barplots showing the proportion of co-expressed STIRs (purple) or control sequences (beige and orange) for the G4NG4 motifs (or their reverse complement), for co-expression in HepG2 cells. Shown are Bonferroni-corrected proportion test P-values. (D) Histogram showing the binned distribution of G-rich motifs (or their reverse complement) across the co-expressed STIRs of HepG2 cells. (E) Metagene analysis (top) and corresponding binding heatmap (center) of MAZ in ENCODE ChIP-seq data in HepG2 co-expressed STIRs and flanking 5’ and 3’ regions and at the control regions. Top graph shows median and standard error and the bottom heatmap shows the binned Bonferroni-corrected paired Wilcoxon rank-sum test P-value heatmap. (F) As in (E), but for the SP1 ChIP-seq data in HepG2 cells.

Interestingly, the enrichments are evident when we consider the G-rich motifs but not necessarily their C-rich reverse complements, pointing to the specific importance of Gs on the transcribed strand (**Fig. 2C-D**), and consistent with the overall enrichment of Gs mentioned above.

MAZ elements are G-rich motifs that were previously associated with efficient termination and are being used as termination signals *in vitro* and *in vivo* at individual genes (Yonaha and Proudfoot 1999; Ashfield, Enriquez-Harris, and Proudfoot 1991a; Ashfield et al. 1994). Since our candidate motifs resemble the ‘MAZ’ element G5AG5, and due to the general enrichment of G-rich motifs, we checked whether there is preferential binding of MAZ in STIRs. Notably, ENCODE ChIP-seq data of MAZ in HepG2 cells displayed a higher binding signal in STIRs compared to both types of controls (**Fig. 2E**). Notably, the majority of the signal stemmed from the promoter of the downstream gene and not the 3’ of the upstream gene or the control. A similar known G-rich element is the Sp1 binding motif (G/T)GGGCGG(G/A)(G/A)(C/T) (Nagaoka, Shiraishi, and Sugiura 2001), and it has been previously proposed that MAZ but not Sp1 contribute to Pol2 pausing downstream of the TTS (Yonaha and Proudfoot 1999). Interestingly, Sp1 also showed a slightly higher binding signal at the promoter regions of STIRs in HepG2 and to a lesser extent in K562 cells (data not shown), but not upstream of the promoter (**Fig. 2F**).

STIRs therefore preferentially harbour G-rich motifs, and those are typically located proximally to the downstream promoter, and are associated with a slightly increased binding of MAZ and to a lesser extent SP1 near the downstream promoter.

### Specific proteins preferentially bind STIRs

These findings led us to seek genome-wide evidence of preferential binding of other proteins within STIRs. Using ENCODE ChIP-seq data, we found enriched binding of several factors, including AGO2, AGO1, RBM22, BCL3, MYNN, and NFATC1 (**Fig. 3A-G** and **S3A-D**). Interestingly, G-rich stretches and AGO proteins were previously implicated in transcriptional termination at specific genes (Skourti-Stathaki, Kamieniarz-Gdula, and Proudfoot 2014). The model of Pol2 pausing following PAS transcription suggests that G-rich terminator elements found tens of bp downstream of the PAS signal further enhance Pol2 pausing. This pausing was suggested to be facilitated by R-loop structures reported to be particularly enriched downstream of the TTS of some genes. A study by (Skourti-Stathaki, Proudfoot, and Gromak 2011) suggested that R-loops at G-rich regions in termination regions of the β-actin gene induce antisense transcription, which leads to the generation of dsRNA and the recruitment of the RNA-interference (RNAi) factors such as AGO1, AGO2, DICER, and the G9a histone lysine methyltransferase. This recruitment was reported to lead to deposition of the H3K9me2 repressive mark and to the recruitment of heterochromatin protein 1γ (HP1γ), which in turn may reinforce Pol2 pausing and promote transcription termination. Since we observed significant binding of both AGO1 and AGO2 – components of the RNAi machinery in STIRs, we sought to further examine the other factors tied with the same pathway in these regions. Analyzing data of RNA:DNA hybrids profiled using the S9.6 antibody in K562 cell line by (Sanz et al. 2016), we indeed observed a substantial enrichment of R-loops in STIRs (**Fig. 4A**). To further examine the model suggested by Skourti-Stathaki *et al*., we examined the binding of the other factors using available ENCODE data, including G9a (EHMT2) – an H3K9 methyltransferase, HP1γ (CBX3), and the H3K9me2 chromatin mark. All these factors were not enriched in STIRs (**Fig. S4A-C**). Conversely, G9a showed ∼30% depletion within STIRs, peaking at the promoter, along with slightly lower levels of H3K9me2 binding overall, precluding the promoter region, which was similarly depleted of H3K9me2 (**Fig. S4A and S4C**). Notably, G9a has additional substrates, such as H3K9 (Shinkai and Tachibana 2011), therefore it might be recruited there under other circumstances. Alternatively, we hypothesized that the relatively low levels of H3K9me2 and its histone lysine methyltransferase within STIRs can be explained by the tendency of tandem genes to reside within relatively gene-rich regions, which are generally less associated with heterochromatic marks (Sanz et al. 2016; Gilbert et al. 2004). Therefore, we next examined the binding of PHF8, an H3K9me2 demethylase (Zhu et al. 2010). Interestingly, PHF8 levels were somewhat higher in control promoter regions than in STIRs, perhaps because it is recruited by H3K9me2, which was also higher in control promoter regions (**Fig. S4C-D**). We conclude that whereas AGO binding and R-loops are prevalent in STIRs genome-wide, there is no genome-wide evidence for preferential activity of the H3K9me2-associated machinery in these regions.

**Figure 3.**
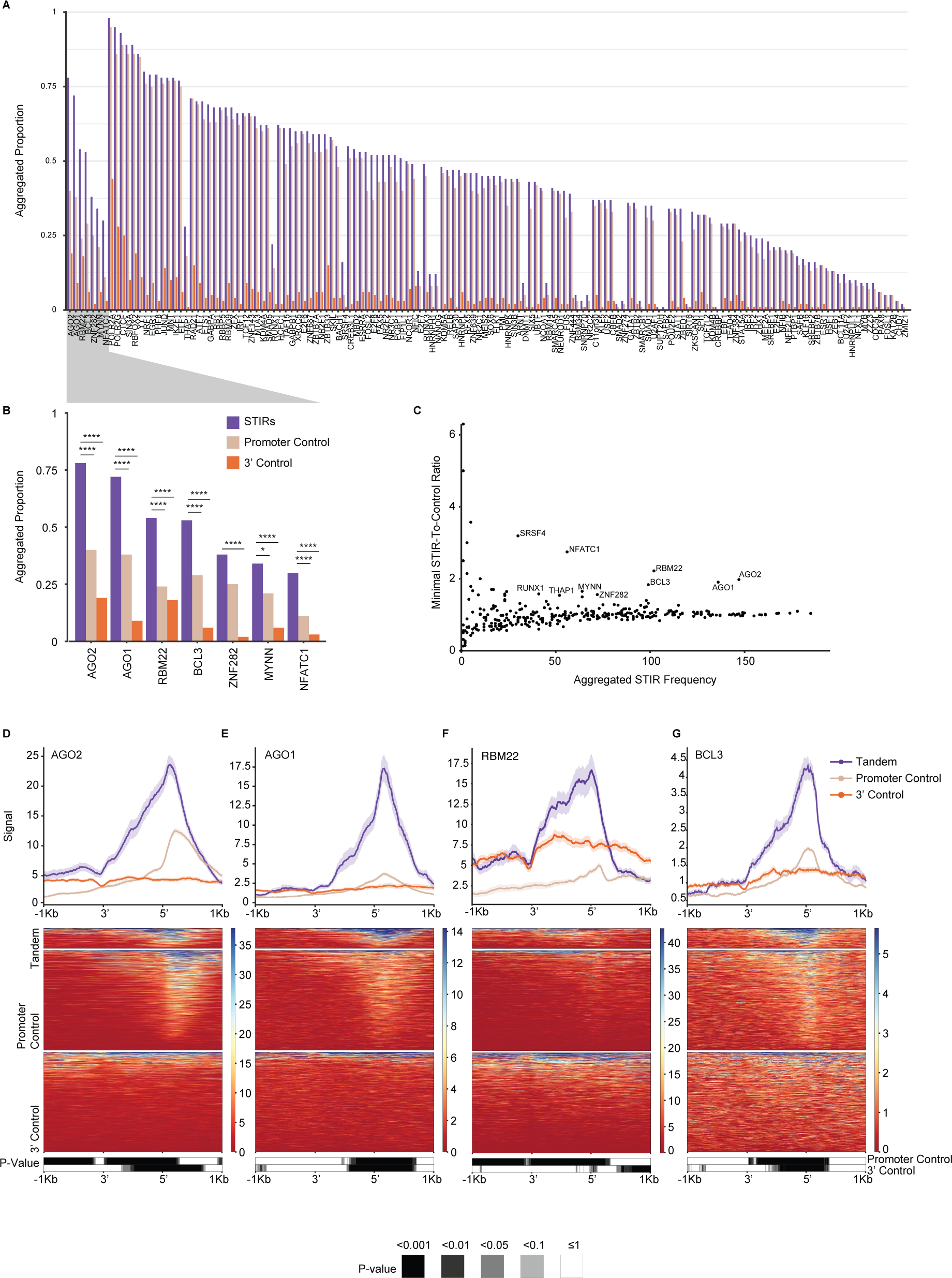
Enriched binding of proteins at STIRs. (A) Barplot showing the proportion of co-expressed STIRs (purple), promoter control- (beige), and 3’ control- (orange) sequences bound by the different proteins (analyzed in HepG2 cells). Bound sequences are aggregated and counted once when multiple binding peaks per sequence were observed. Shown are only proteins with higher binding frequency at STIRs over both controls. Proteins are ordered by ranked frequency in STIRs and ranked descending order of calculated minimal ratio of frequencies between STIR and each of the two controls. Shown are Bonferroni corrected proportion test P-values (*:P<=0.05, **:P<=0.01, ***:P<=0.001, ****<=0.0001). P-values for the equivalent proportions in K562 cells are similar except for MYNN promoter control versus STIR P=2×10^-3^ (**) (B) Zoom-in on the top STIRs HepG2 protein binding candidates. (C) Scatter plot showing the number of HepG2 co-expressed STIRs bound by each protein (as in (A)) versus the minimal tandem-to-control ratio calculated for each control. Indicated are the top proteins enriched in STIRs. Y-axis values>6 were filtered out. (D-G) Metagene analysis (top) and corresponding binding heatmap (center) of median ChIP-seq signal of selected STIR-binding protein candidates AGO2 (D), AGO1 (E), RBM22 (F) and BCL3 (G). Bottom heatmap corresponds to binned paired Wilcoxon rank-sum tests (Bonferroni corrected).

**Figure 4.**
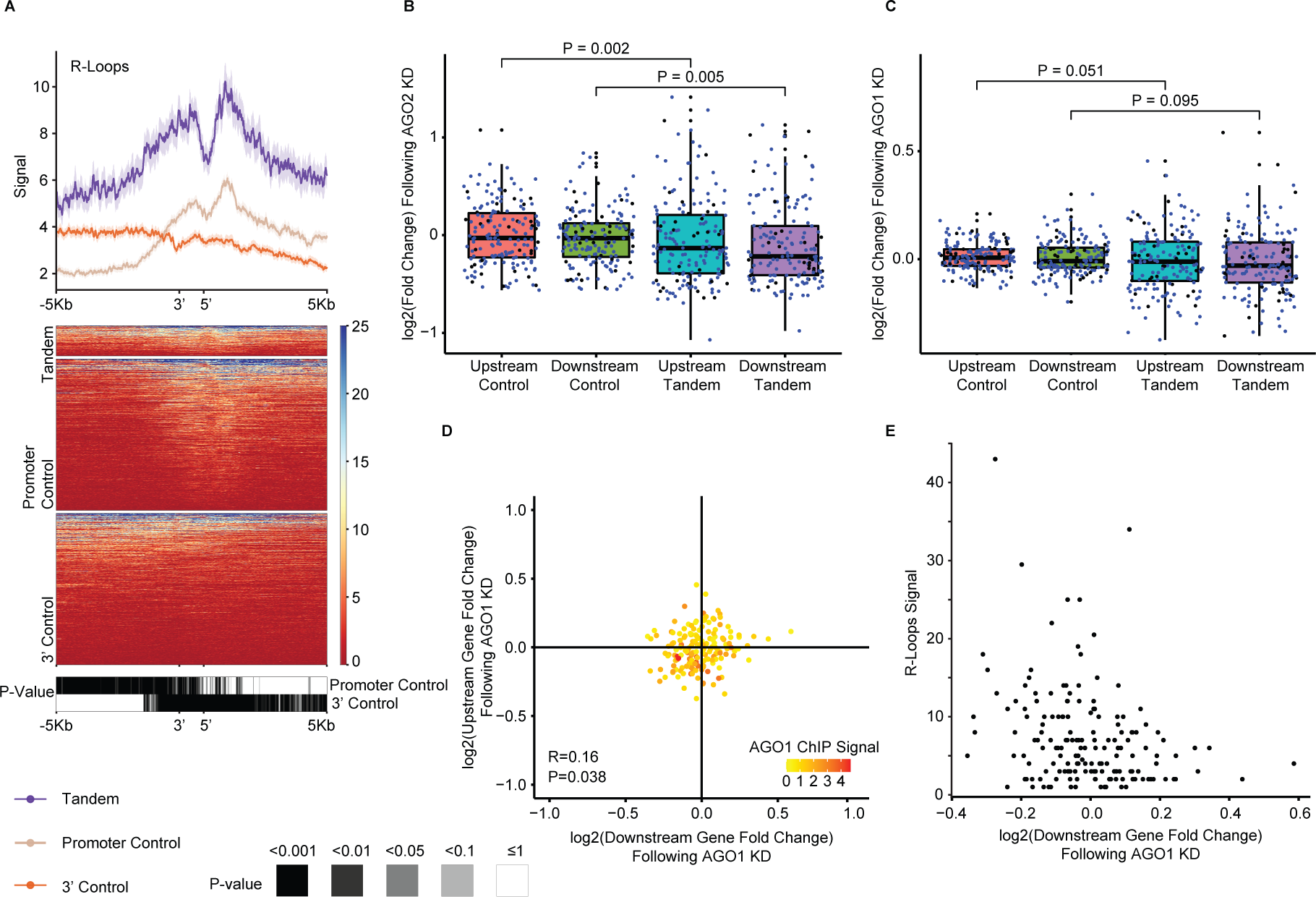
Features promoting the proper expression of tandem gene pairs. (A) Metagene analysis (top) and corresponding heatmap (center) showing S9.6 DNA:RNA antibody DRIP-seq median signal in K562 co-expressed STIRs and flanking regions and in their controls. Bottom heatmap shows the corrected P-value of binned paired Wilcoxon rank-sum test. Data from Sanz *et al*., 2016. (B) Boxplot of expression changes in upstream or downstream co-expressed tandem genes following AGO2 KD in K562 cells (Data from ENCODE). Five genes were used as controls per tandem gene (see Methods, **Fig. S1**) and aggregated using the average log2-transformed fold change value of each quintet. Blue dots correspond to tandem pairs co-expressed in both K562 and HepG2 cell lines (or their respective control). Black dots are tandem genes co-expressed only in the respective cell line. P-values were obtained using paired Wilcoxon rank-sum tests. (C) As in (B), for AGO1 KD in K562 cells. (D) Scatter plot showing the changes in expression following AGO1 KD in K562 cells. Each dot represents the changes in expression of a single tandem pair. Colors indicate median AGO1 ChIP-seq signal per STIR. Calculated Pearson’s correlation coefficients between upstream- and downstream-tandem genes changes in expression and P-value are indicated. Spearman correlation between the downstream- or upstream-gene expression changes following KD and median AGO1 ChIP-seq signal at STIR was tested, with coefficients of -0.134 or -0.295 and P=0.088 or P=1.2×10^-4^, respectively. Wilcoxon test of STIR ChIP-seq signal tested between the third quadrant and all other quadrants: P=1.7×10^-3^. (E) Scatter plot showing median R-loop signal at STIRs as a function of the changes in expression of the downstream gene in the co-expressed tandem pairs following AGO1 KD in K562 cell line. Spearman correlation coefficient of –0.278, P=3.1×10^−4^.

### AGO2 binding promotes the expression of tandem gene pairs

Notable enrichment of AGO1 and AGO2 binding in STIRs led us to further look for possible consequences of the binding. We examined the changes in the ENCODE gene expression dataset of polyadenylated RNA following AGO1 and AGO2 knockdown (KD) in K562 and HepG2 cell lines. For AGO2 KD we observed a significant yet mild decrease in the expression of the tandem co-expressed genes compared to their respective controls (**Fig. 4B**). For AGO1, we observed smaller changes in the same direction, which were significant only in HepG2 cells, and only for the downstream gene in the tandem pair (**Fig. 4C** and **S4E**). Examination of the correlation between the downstream and upstream tandem genes expression changes following the KD showed little to no correlation between the tandem pairs (**Fig. 4D** **and S4G-H**), except the notable scarcity of tandem gene pairs where both genes were up-regulated following AGO2 KD (**Fig. S4H**).

While there was no significant change in expression when considering all the tandem pairs, when we integrated ENCODE K562 AGO1 ChIP-seq data, we observed increased binding of AGO1 to chromatin in the STIRs that were associated with relative downregulation of gene expression of both the upstream- and downstream-genes following the KD (Wilcoxon test P<0.05). Furthermore, we found a significant negative correlation between AGO1 binding and the change in the expression of the upstream gene following KD (Spearman’s correlation coefficient: –0.295, P=1.2×10^−4^) and a similar yet non-significant (Spearman’s correlation coefficient: –0.134, P=0.088) effect for AGO1 binding and the change in expression of the downstream gene (**Fig. 4D**). In addition, examining the connection between R-loop signal and AGO1 KD in K562 cells showed significant correlation between changes in the downstream co-expressed tandem gene expression and R-loop signal (P=3.1×10^−4^) (**Fig. 4E**). However, R-loop signal intensity did not correlate with gene expression changes of co-expressed tandem genes following AGO2 KD in K562 cells (**Fig. S4H**).

### Pol2 marked with specific modifications accumulates in STIRs

Due to the short distance between the tandem gene pairs that we considered, and the continued association of Pol2 with DNA downstream of the CPA site, with the addition of our constraints for picking only pairs of genes that co-express in the relevant cell type, we reasoned that STIRs might be associated with distinct patterns of Pol2 occupancy. To examine that, we used mammalian Native Elongation Transcript sequencing (mNET-seq) data from (Schlackow et al. 2017). The mNET-seq strategy uses antibodies for several different Pol2 CTD marks, to obtain segments of nascent-RNA bound by Pol2 in its different states. Importantly, in contrast to ChIP-seq, mNET-seq data are strand-specific, allowing to consider Pol2 traveling strictly on the same strand as the considered tandem genes. We used the same definitions described above to define a set of 159 tandem genes co-expressed in HeLa cells and their controls. Overall, ∼1.25 fold enrichment over the promoter control of total Pol2 (phosphorylated and non-phosphorylated, tested using CMA601 antibody) was observed at the peak just downstream of the promoter in the STIRs (**Fig. 5A** and **S5A**). There was also substantial Pol2 presence within the STIR whereas it was absent from both controls (**Fig. 5A** and **S5A**), resulting in an overall 6.8 and 8.4-fold enrichment of Pol2 occupancy across the STIR, for the 3’ control and promoter control, respectively (see Methods). Interestingly, all modifications of Pol2 showed enrichment at tandem intergenic regions and at the promoters with slightly different patterns. For example, T4P modification (using 6D7 anti-CTD Thr4-P modification antibody) showed a stable signal for the 3’ control, downstream of the TTS, and a gradually increasing signal that was enriched by ∼7 fold compared to the 3’ control in STIRs (**Fig. 5B**). Surprisingly, the signal peaked at approximately the promoter of the downstream tandem gene followed by a decreasing signal in the following 0.5kb that eventually reached background levels. As expected, T4P signal was generally depleted at the promoter controls (**Fig. 5B** and **S5G**). Another notable example is the S2P modification (tested with CMA602 Ser2-P CTD antibody), which showed an enrichment along the STIR that peaked downstream of the transcription start site and was then almost completely absent after 1kb. Notably, in both controls, S2P seemed to be depleted, with a low signal at the promoter control just downstream of the promoter (**Fig. 5C** and **S5B**). Y1P modification was reported to be essential for the function of Positive Transcription Elongation Factor b (P-TEFb) in phosphorylating Ser2 and transitioning from transcription initiation to elongation (Mayfield et al. 2019), S5P modification was suggested to facilitate the recruitment of the stabilizing capping enzyme complex (Ho and Shuman 1999), and S7P is enriched at promoters and gene bodies, and is thought to regulate snRNA biogenesis (Egloff et al. 2007; Harlen and Churchman 2017). For these three modifications, both the tandem genes and the promoter control show a peak of signal enrichment just downstream of the promoter, with ∼1.25-, ∼4-, or ∼2.5-fold higher signal at tandem downstream promoter over the promoter control genes, for Y1P, S5P or S7P, respectively (tested using 3D12 Tyr1-P CTD antibody, CMA603 Ser5-P CTD antibody, and 4E12 Ser7-P antibody, respectively). Interestingly, these signals exist also within the STIR, whereas they are depleted upstream of the promoter controls (**Fig. 5D-F** and **S5C-D,F**). Intriguingly, treatment with Pladienolide B (Pla-B), a splicing inhibitor that binds SF3B1 spliceosome subunit, made the pattern of S5P Pol2 binding almost indistinguishable between the tandem genes and the promoter controls, suggesting that splicing substantially contributes to the Pol2 accumulation in the intergenic region and around the promoter of the downstream gene. Potentially, the inhibition of splicing affects elongation of Pol2 through the upstream gene, leading to paucity of Pol2 around gene end, which is difficult to measure for 3’ control regions, since Pol2 levels near gene ends are generally very low. Alternatively, Pol2 that is stalled at the intergenic region is associated with the transcripts produced at the transcription round that just terminated but are still engaged in the midst of splicing, and so splicing inhibition causes disengagement of such immobilized Pol2 (**Fig. 5G** and **S5E**). Interestingly, the overall presence of Pol2 within STIRs is dramatically higher than in the 3’ parts of transcribed genes, where the occupancy of Pol2, including all the different modifications, is barely over background levels (**Fig. 5A-F**).

**Figure 5.**
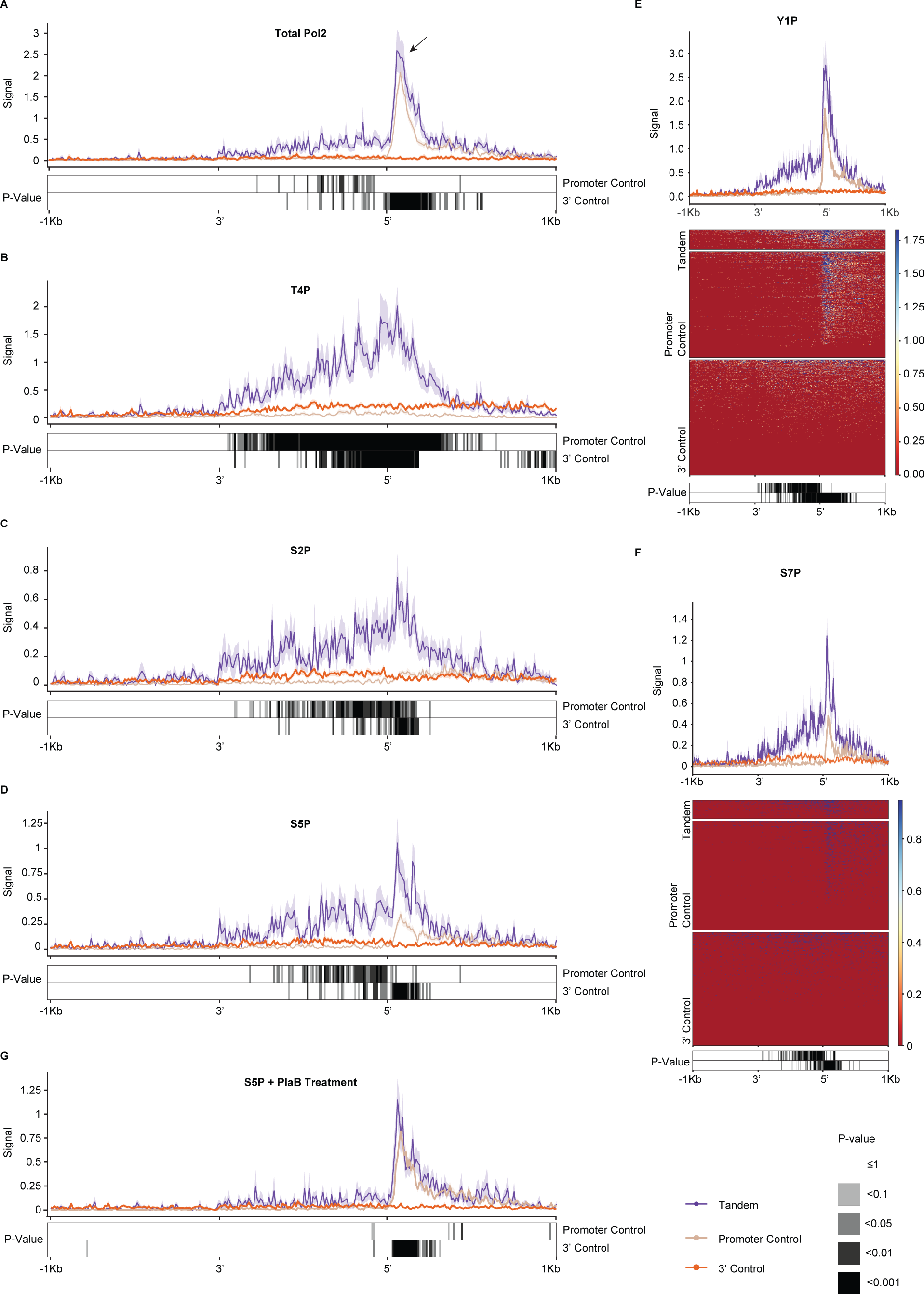
Pol2 accumulates in STIRs. (A-D,G) Metagene analysis (top) showing the different median Pol2 carboxy-terminal domain modifications (phosphorylated and non-phosphorylated, T4P, S2P, S5P and S5P+Pla-B, respectively) occupancy signal at the sense strand of STIRs or controls and their flanking regions. Bottom are the respective corrected paired Wilcoxon rank-sum test P-values. Data from Schlackow *et al*., 2017. (E,F) Metagene analysis (top) and corresponding heatmap (center) of median Pol2 CTD modifications (Y1P and S7P, respectively) mNET-seq signal and standard error at STIRs and controls, and their flanking regions. Bottom are the respective corrected Wilcoxon-paired test P-values. Data from Schlackow *et al*., 2017.

As further evaluation of the role of upstream gene elongation in the accumulation of Pol2 in STIRs, we also examined the consequences of loss of Spt6 and Rtf1, two factors reported to separately control Pol2 progression. We used mNET-seq data from K562 following Spt6 and Rtf1 depletion from (Žumer et al. 2021) and examined Pol2 distribution within STIRs (**Fig. S6**). Spt6 was shown to assist progression of elongating Pol2 through nucleosomes, and its depletion was shown to have little to no effect on Pol2 accumulated at promoters (Žumer et al. 2021). Indeed, data in K562 recapitulated the pattern we observed in HeLa cells, showing strong accumulation of Pol2 in STIRs in control conditions (DMSO-treated cells), which was strongly reduced upon targeted degradation of Spt6 (**Fig. S6A-B**), with little to no effect on Pol2 in the downstream promoter. Interestingly, no effect was seen for Rtf1 depletion (**Fig. S6C-D**).

### NELF-E KD suggests involvement in transcription regulation of downstream tandem genes

We next tested whether the expression of tandem genes is particularly sensitive to the loss of specific protein factors. We examined this by analyzing KD data of 245 different protein factors, in 440 experiments (in HepG2 or K562 cell lines). For most factors, we found concordant changes in expression of the upstream and downstream genes (**Fig. 6A**), possibly related to their overall shared features (see Discussion), and so we were particularly interested in factors whose loss preferentially affected just the upstream or the downstream gene. This analysis highlighted Negative Elongation Factor Complex Member E (NELF-E) as a candidate for the transcriptional regulation of the downstream gene in a tandem pair. NELF-E is a part of the NELF complex (composed of units A,B,C/D and E). Based on current knowledge, release of Pol2 from the proximal-promoter region and its conversion to elongating state is dependent on the displacement of the negative elongation factor (NELF) complex from the nascent transcript. This process is thought to include the phosphorylation of several factors including Ser2 of the CTD, NELF-E, and the Spt5 subunit of the DSIF (DRB Sensitivity Inducing Factor) complex by positive transcription elongation factor b (P-TEFb). These phosphorylations are followed by NELF dissociation and Pol2 transition from promoter-proximal pausing to elongation (Lu et al. 2016). NELF-E KD in HepG2 cells led to the greatest median fold change in expression of the downstream tandem gene versus its controls accompanied by a negligible median KD effect over the upstream tandem gene compared to its controls (**Fig. 6B** and **Table S10**). The trend was similar in K562 cells yet the effect was less robust (**Fig. S7A**). Interestingly, NELF-E ChIP-seq data in K562 cells showed slightly lower signal at the downstream tandem promoters relative to controls (**Fig. S7B**). In addition to NELF-E, we found SMN1 and XRN2 KD as having similar, yet less robust effects over the downstream and not the upstream tandem gene (**Fig. 6A** and **S7C-D** and data not shown). Intriguingly, SMN was previously suggested to play a role in R-loops resolution (Jangi et al. 2017; Zhao et al. 2016). These results suggest that NELF-mediated Pol2 pausing preferentially restricts expression of downstream genes found in tandem pairs, which is potentially related to the accumulation of Pol2 within STIRs and over the promoters of the downstream genes.

**Figure 6.**
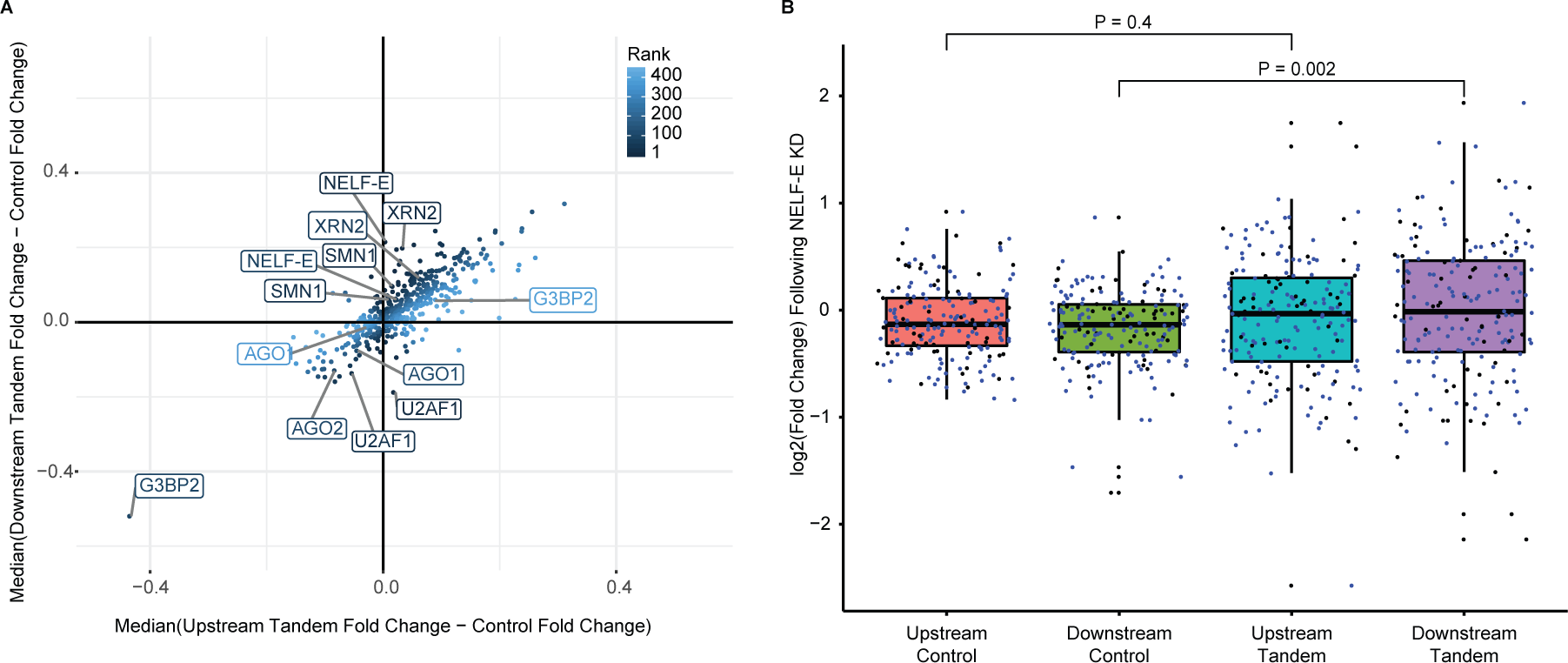
NELF-E involvement in transcription regulation of downstream tandem genes. (A) Scatter plot showing the median difference in expression changes between downstream tandem (y-axis) or upstream tandem (x-axis) genes and the averaged expression changes in control. Candidate genes with high KD effect on expression compared to controls in the downstream tandem gene but not in the upstream tandem gene are ranked lower and appear darker and vice versa. Shown are results from 440 KD experiments done in either HepG2 or K562. KDs with the lowest ranking, and other proteins of interest are marked. (B) Boxplot of expression changes in upstream or downstream co-expressed tandem genes following NELF-E KD in HepG2 cells (data from ENCODE) or their averaged aggregated controls. Blue dots correspond to tandem pairs co-expressed in both K562 and HepG2 cell lines (or their respective control). Black dots are tandem genes co-expressed only in the respective cell line. P-values were obtained using paired Wilcoxon rank-sum tests.

## Discussion

Our results show that STIRs differ from regions flanking 5’ or 3’ ends of other genes in sequence composition, protein associations and in the presence of Pol2 associated with different CTD modifications. These regions primarily resemble promoters, which implies that sequences required for regulated transcription initiation are longer and/or experience a stronger selective pressure compared to sequences required for efficient termination, and yet differ from other promoters, in particular in strong accumulation of Pol2 bearing CTD modifications traditionally associated with gene ends.

Our observation of a slightly elevated Guanine content at the tandem intergenic regions was accompanied by enrichment of specific G-rich motifs in these regions compared to the control regions. As mentioned above, these findings echo studies of individual genes showing several possible functions for G-rich sequences. These past reports include demonstrations of links between the G-rich regions and MAZ, and SP1 (Dalziel, Nunes, and Furger 2007; Arhin et al. 2002; Oberg et al. 2005), and with the formation of R-loops. GGGGAGGGG “MAZ” elements positioned in proximity to the polyA site was reported to contribute to Pol2 pausing and thus promote more efficient transcription termination facilitated by the XRN2 exonuclease. Interestingly, MAZ itself does not seem to have an effect on the termination activity, as RNAi-mediated depletion of MAZ did not affect termination (Gromak, West, and Proudfoot 2006). This observation fits our analysis showing lack of enrichment of MAZ binding at either 3’ control TTS region and the TTS of the upstream gene in a tandem pair (**Fig. 2F**) despite the overall G-richness of the tandem intergenic regions (**Fig. 1D-E**). These findings together may point towards the importance of the G-rich structure of the sequences themselves, e.g., for regulation of Pol2 speed, or to binding of other protein(s), rather than MAZ. Importantly, Gromak *et al*. reported that the proximity of MAZ elements to the polyA site is important for the termination process, as a construct containing MAZ element at a 2kb distance from the polyA site, rather than a proximal one, significantly reduced the termination efficiency. Our analysis only partly supports this notion, since we observed no bias for the MAZ elements to appear in proximity to the TTS (**Fig. 2F**). MAZ elements and not GGGGCGGGG “SP1 elements” were proposed to specifically promote Pol2 pausing and polyadenylation (Yonaha and Proudfoot 1999). Interestingly, the tandem intergenic regions we analyzed were enriched with ‘GGGGCGGG’ or ‘GGGGCGGGGSC’ motif, that resembles the SP1 element rather than the MAZ element (**Fig. 2A and 2C-c**). Furthermore, all types of G-rich motifs (with the notable exception of (G)_9_) were much more prevalent upstream of the promoter controls, rather than at 3’ controls, and indeed we observed binding of both SP1 and MAZ centered at promoters rather than at the 3’ ends. Our findings thus do not support the notion of the importance of specifically an Adenosines flanked by runs of Guanines in facilitating Pol2 pausing at termination sites (**Fig. 2D**).

Dense genes are associated with shorter introns (Amit et al. 2012), which fits our observation of shorter total intronal length in tandem genes as a group (**Fig. 1C**). Other features associated with shorter introns include lower levels of low complexity sequences, higher GC-content, proximity to another transcription unit and slower transcription rate (Veloso et al. 2014). Intriguingly, most of these features are exhibited to some extent at the tandem genes we analyzed, in some cases within the intergenic region rather than in the gene bodies, such as the lower rate of transposable elements (**Fig. 1F**) and the higher GC-content (**Fig. 1D**). Transposable elements are the main source for intronal expansion ((Wu et al. 2013)). The lower rate of transposable elements at STIRs is potentially related to the enrichment of Pol2 (**Fig. 5**), which may provide less opportunity for their introduction to these regions. Interestingly, we also find an overall higher level of sequence evolution in STIRs compared to control regions (**Fig. 1G**). This increase is possibly related to the potential of the G-rich motifs to form G-quadruplex structures, which are associated with genome instability and increased mutation rate (Bochman, Paeschke, and Zakian 2012).

During transcription termination, the current model is that transcription continues for a few hundreds to several kbs post the TTS before the polymerase is released from the DNA, accompanied by the T4P CTD modification. Therefore, it is unexpected that we find extensive Pol2 signal in the intergenic region of tandem genes in the “total Pol2” mNET-seq data (using anti-CTD pan-Pol2 CMA601 antibody) and when examining Pol2 with all the different CTD modifications, including high levels of T4P signal downstream of the promoter while being almost at background levels in both controls. Particularly striking is the prominent peak of T4P modified Pol2 at the downstream promoter, whereas such Pol2 is rarely found at control promoters. One potential explanation for this observation is that there is a unique regulatory regime taking place when Pol2 is transcribing tandem transcriptional units. In that case, it is possible that the STIR may constitute a “preparation area” for a new cycle of transcription of the downstream tandem genes. This would include slowing down of Pol2 at the intergenic region, which may be supported by its enriched signal at 3’ ends of upstream tandem genes over control genes, where Pol2 is not necessarily recycled or where recycling is distributed over a substantially longer genomic sequence, or where fast release of Pol2 might be potentially favored, to maintain its nuclear pool. In tandem genes, the Pol2 that finished transcribing the upstream gene can potentially be used to transcribe the downstream one, perhaps before its CTD marks ‘reset’ from their termination-associated state.

In order to examine which region of the STIR constitutes the primary accumulation site of Pol2, we examined Pol2 occupancy in tandem intergenic regions of increasing sizes (while keeping the other criteria the same, **Fig. S8**). We observed that with increasing intergenic distances there was a less pronounced Pol2 accumulation in the intergenic region, and that Pol2 accumulation was predominantly found near the downstream promoter. This suggests that Pol2 that finishes the transcription of the upstream gene continues to be associated with chromatin and predominantly pauses near the downstream promoter (while carrying the termination-associated marks T4P and S2P). Notably, this pausing is potentially facilitated by the G-rich sequences that are also predominantly found near the downstream promoter (**Fig. 2E**). Furthermore, our finding that NELF-E depletion preferentially leads to an increase in expression of the downstream genes in the tandem pairs suggests that the Pol2 pausing may have a functional role in restricting expression from the downstream promoter.

As mentioned above, additional features of tandem genes may also support slower Pol2 dynamics within STIRs. Alternatively, it is also plausible that although we only considered tandem genes that are co-expressed in bulk RNA-seq data, the actual transcription cycles of the two tandem genes are disjoint events. This may be supported by the T4P signal we see at the promoter region. The signal may stem from Pol2 which has not yet dissociated from the DNA after transcribing the upstream gene, and is not going to be involved in the transcription of the downstream gene. If this is the case, since we controlled for expression, we would expect the STIR profile to resemble the superposition of the signals of both types of controls. Notably, this does not appear to be the case, at least when considering the median Pol2 occupancy signal.

When considering chromatin marks in STIRs, as mentioned above, we did not observe a notable enrichment of H3K9me2 in STIRs, where it was depleted similarly to other promoters. Other histone marks also showed largely unremarkable patterns within STIRs (**Fig. S9**), which were largely superpositions of the patterns of the control regions, with a notable exception of H3K79me2, which showed a reduced pattern in the gene body of the downstream gene. For H3K36me3, the reduction to background levels was faster in STIRs compared to the 3’ controls, likely a consequence of the nucleosome-depleted region at the downstream promoter. Similarly, for H3K4me3, we found a smaller and narrower peak centered at the -1 nucleosome, likely reflecting reduced levels of divergent transcription from the downstream gene promoter.

Finally, we considered the possibility that different gene subsets are responsible for the enriched binding patterns of the various factors that we found enriched within STIRs, and/or that different factors tend to preferentially co-bind the same regions. To test this, we clustered the binding data (**Fig. S10-S11** and **Tables S11-S12**), however clustering of both the STIRs and the factors did not point toward a specific regulatory pathway or subsets of tandem genes with the same protein binding patterns.

An immense amount of research has been dedicated so far into understanding transcription initiation in mammalian cells, and relatively less attention has been dedicated to transcriptional elongation and termination. Still, these events were usually studied in isolation, e.g., by considering separately promoters and termination regions. Our results suggest that the presence of an upstream termination region within up to 2kb can have a dramatic effect on the protein occupancy at promoters, and those proteins lead to a significant effect on the transcriptional activity of the downstream genes. We further found that some of the features previously associated with efficient termination at individual genes (e.g., G-rich elements and MAZ binding) are likely mostly related to downstream promoters and not the upstream genes. Together with the emerging importance and understanding of architectural chromatin domains, this further suggests that integrative analysis of gene regulation on the genome rather than on the single-gene level will most likely be required for detailed and accurate models of gene regulation.

## Methods

### Extraction of tandem genes

46,012 Human hg19 RefSeq coding transcript annotations were downloaded from UCSC Genome Browser (GB). Unique start and end data per gene were kept to remove different inner splicing variants. Non-coding transcripts of these genes were integrated and multiple transcripts of the same gene were flattened, choosing the minimal start coordinate and maximal end coordinate. Genes from the 23 aligned chromosomes were kept leaving 19,464 flattened transcripts. Extraction of the 5 closest downstream (for the definition of tandem or convergent genes) or upstream (for the definition of divergent genes) genes per gene was done using the ‘closest’ method of the bedtools-gnu/2.25.0 package. Pairs of genes were filtered to keep only the non-overlapping ones, and only the ones where both genes are longer than 5kb and shorter than 800kb. Genes of the different orientations were then divided into 4 groups based on their minimal distance from one another, as reported by bedtools.

### Mouse homologs analysis

Mouse Refseq curated mm10 genes were downloaded from GB and were processed in a similar manner to the human genes to obtain 497 mouse tandem genes. Mouse genes with homology to the human tandem gene were obtained using the Ensembl database (Kinsella et al. 2011), and mouse pairs were kept if both were tandem in both mouse and human and marked as having “one2one” orthology type.

### Determining co-expressed tandem gene set

ENCODE expression data for multiple cell lines was quantified using RSEM as described (Zuckerman and Ulitsky 2019). The tandem pairs were first filtered for expressed pairs (requiring both genes with expression >2TPM). Differences between the upstream and downstream tandem genes expression level were calculated, transformed to absolute values and empirical cumulative distribution function was operated over the values. The 75^th^ percentile was chosen as the maximal allowed expression difference threshold between the two co-expressed genes (for example 66.37 and 82.12 for HepG2 and K562, for the set of tandem genes), leading to 188 and 164 co-expressed tandem pairs in HepG2 and K562, respectively.

### Creating the control set for the co-expressed tandem genes

A set of 8,861 non-tandem genes was defined as genes >5kb and <800KB with minimal distance >5kb from another gene on any strand. Total intronal length was calculated for both tandem and non-tandem genes based on the average total intronal length of all the isoforms of that gene annotated by Refseq. Each co-expressed tandem gene was paired with 5 control genes with the closest weighted resemblance to it both in expression levels and in total intronic length (For example, for the HepG2 co-expressed pair MRM2 and MAD1L1 which have TPMs of 12.68 and 14.02 and total mean intronal length of 6,307 and ∼366kb, respectively, respective control genes were RRP7A with TPM of 12.33 and intronal length of 6,034 and ASAP1 with TPM of 12.94 and intronal length of ∼385kb. For both types of controls, sequences with the same length as the length of the intergenic region of the original tandem pair were extracted either upstream of the promoter region (“promoter controls”, controlling for the downstream co-expressed tandem gene) and downstream of the 3’ end (“3’ control”, controlling for the upstream co-expressed tandem gene) and were set as the control set for the intergenic region of the tandem pair.

### Sequence analysis

FASTA format sequences of the intergenic region of HepG2 and K562 co-expressed tandem pairs and their control sequences were extracted. The sequences were divided into ∼70 bins (based on the minimal length of the co-expressed tandem intergenic regions), and the nucleotide composition ratio was calculated. Control sequence compositions were aggregated using the mean of each set of 5 controls. Significance was tested using paired Wilcoxon rank-sum tests. Genomic repeats overlapping individual positions within the tandem intergenic regions and control sequences were obtained usingRepeatMasker tracks of GB and were filtered to keep only non-simple repeat annotations. Bedtools’ maskfasta function was used for soft masking and FASTA sequences for the tandem co-expressed genes and controls were extracted and the sequences were binned as described above. PhyloP evolutionary conservation scores were obtained from GB, and plots were created using the deepTools package (Ramírez et al. 2014).

### Enriched motifs in co-expressed tandem intergenic regions

188 and 164 non-masked tandem intergenic region sequences of HepG2 and K562 co-expressed tandem genes were used as input for STREME which yielded 37 motifs with P<0.05. Motif occurrences were counted in the tandem intergenic region- and control sequences of the cell line they were originally detected at using FIMO with STREME output. Scores were summarized and motifs were filtered to keep ones with higher prevalence in the tandem intergenic regions both in the measure of total number of motif occurrences per sequence and in the relative number of genes carrying the motif relative to the controls. Filtered motifs were ranked by the minimal tandem-to-control scores ratio and ordered by the sum of ranks of both measures. Proportion tests were applied over the filtered motifs and P-value was corrected using Bonferroni correction.

### TF binding and metagene analysis

ChIP-seq binding cluster data across the human genome were obtained for 338 proteins profiled by the ENCODE project from the GB (**Table S13**). Binding sites were intersected with the co-expressed tandem intergenic regions and control sequences of HepG2 or K562 cells using the intersect function of bedtools. Enriched TF binding at co-expressed tandem intergenic regions over controls was calculated by taking the minimal ratio value between the prevalence of TF binding at the tandem regions fractionated by either the promoter- or 3’-control regions normalized binding prevalence. For example: AGO1 binding sites intersected at least once with 136 out of 188 (∼72%) of co-expressed tandem intergenic region sequences, but only 357 or 89 of the 940 (∼38% and ∼9%) of the promoter and 3’ control sequences in HepG2, respectively. The analysis produced a set of 6 TFs with the highest minimal ratio between co-expressed tandem intergenic region- and control-binding which were found in both HepG2 and K562 cell lines. To further examine the binding pattern throughout the tandem intergenic- or control-regions, ChIP-seq data of the 6 candidates was obtained from the ENCODE project and was visualized as a metagene plot using deepTools (Ramírez et al. 2014), averaging the bins using the median value and showing the standard error. In addition, each ChIP-seq experiment was adjusted with its own set of co-expressed tandem genes and control genes based on the specific cell type used for the ChIP-seq experiment as explained previously. Metagene plots for EHMT2, H3K9me2, PHF8 and HP1γ ChIP-seq data (from the ENCODE project) and R-loop data (from the GEO database, accession <u>GSE70189</u>) were drawn in a similar manner.

### Knock-Down analysis

ENCODE shRNA experiments of 245 genes (440 experiments) done in K562 and/or HepG2 were analyzed using DESeq2 (**Table S14**). For each factor, data of the co-expressed tandem genes and their controls within the respective cell line was extracted and the distribution and log2-transformed fold change of gene expression was plotted and tested using Wilcoxon paired rank-sum test for the set of upstream or downstream genes within the co-expressed tandem pairs and their averaged controls. Scatterplot of the changes in expression following KD of AGO1 and AGO2 for the downstream and upstream co-expressed tandem genes were plotted and tested using Pearson’s correlation. Additional correlation tests for ChIP-seq or R-loops median signal and co-expressed tandem genes expression change following KD were done using Spearman’s correlation. Median difference in expression changes following KD between the upstream or downstream gene and the controls were calculated. KD experiments were ranked based on minimal-to-maximal effect over the upstream gene compared to controls, maximal-to-minimal effect over the downstream gene compared to controls, and overall absolute difference between the effect on the upstream and downstream gene. Ranking was done so that KD targets with low ranks are those affecting mostly the downstream tandem genes (and not the upstream tandem genes) and vice versa.

### Pol2 modifications analysis

Data of Pol2 binding was downloaded from the GEO database (accession <u>GSE81662</u>) in bigWig format. Matrices for the plus and minus strand of the co-expressed tandem intergenic regions in HeLaS3 and their respective controls were built using deepTools and binning was done based on the median value. Matrices were then filtered according to strand and were combined. Extreme values were filtered to remove rows containing outliers with values above 99.99%- and under 0.01%- of the total combined matrix. To calculate the overall enrichment of total Pol2 at STIRs over control 3’ or promoter regions, raw output sense of “total Pol2” (GSM2357382) matrices were filtered to keep only the intergenic region, and overall signal was summed at STIRs, or averaged per control quintet of each STIR and then summed.

### Clustering of tandem genes and binding experiments

Maximal value per gene per binding experiment was calculated over the intergenic region and flanking regions (1kb on each side, unless stated otherwise) and over the associated control regions, using the deepTools matrix output. For the control experiment, each quintet controlling for a single tandem gene was first aggregated using mean value to create a mean matrix per experiment. Tandem max matrix was divided by either control matrix, genes with over 17 missing values across experiments were removed from the cluster analysis, matrix values exceeding 10 were converted to 10. Clustering of the columns was done using Pearson’s correlation using the “complete.obs” option to handle NA values and complete-link measure. Genes were clustered using complete-link measure and euclidean distances method. Log2-transformed FPKM values for gene expression annotations were computed using the ENCODE RNA-seq data.

## Supplementary figures

**Figure S1.**
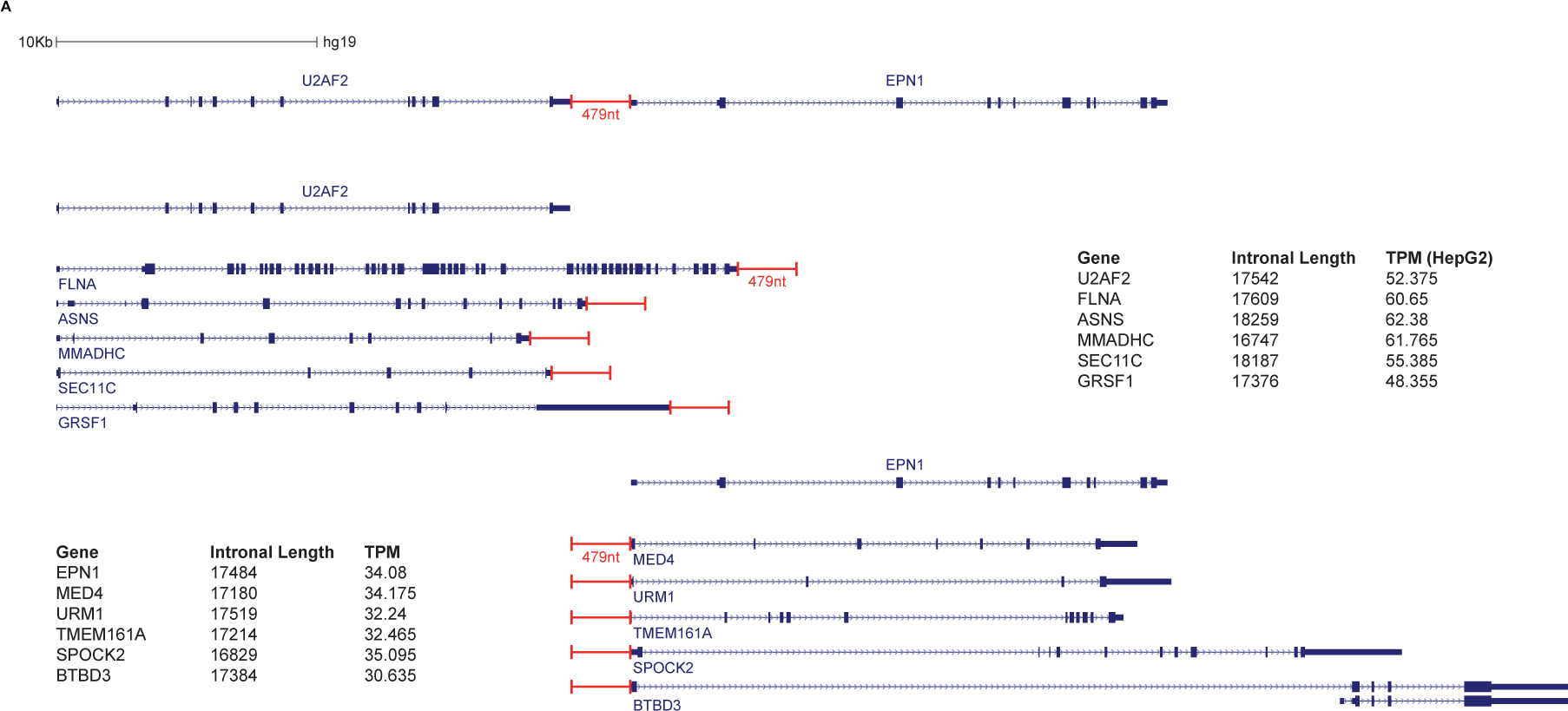
Definition of controls. (A) An example of the set of controls for the U2AF2-EPN1 tandem pair separated by 479 nucleotides (top). U2AF2 upstream tandem gene was matched with five 3’ controls having the closest total intron length (averaged over the different RefSeq transcripts) combined with similar expression (center). EPN1 downstream tandem gene was matched with five promoter controls in the same manner (bottom).

**Figure S2.**
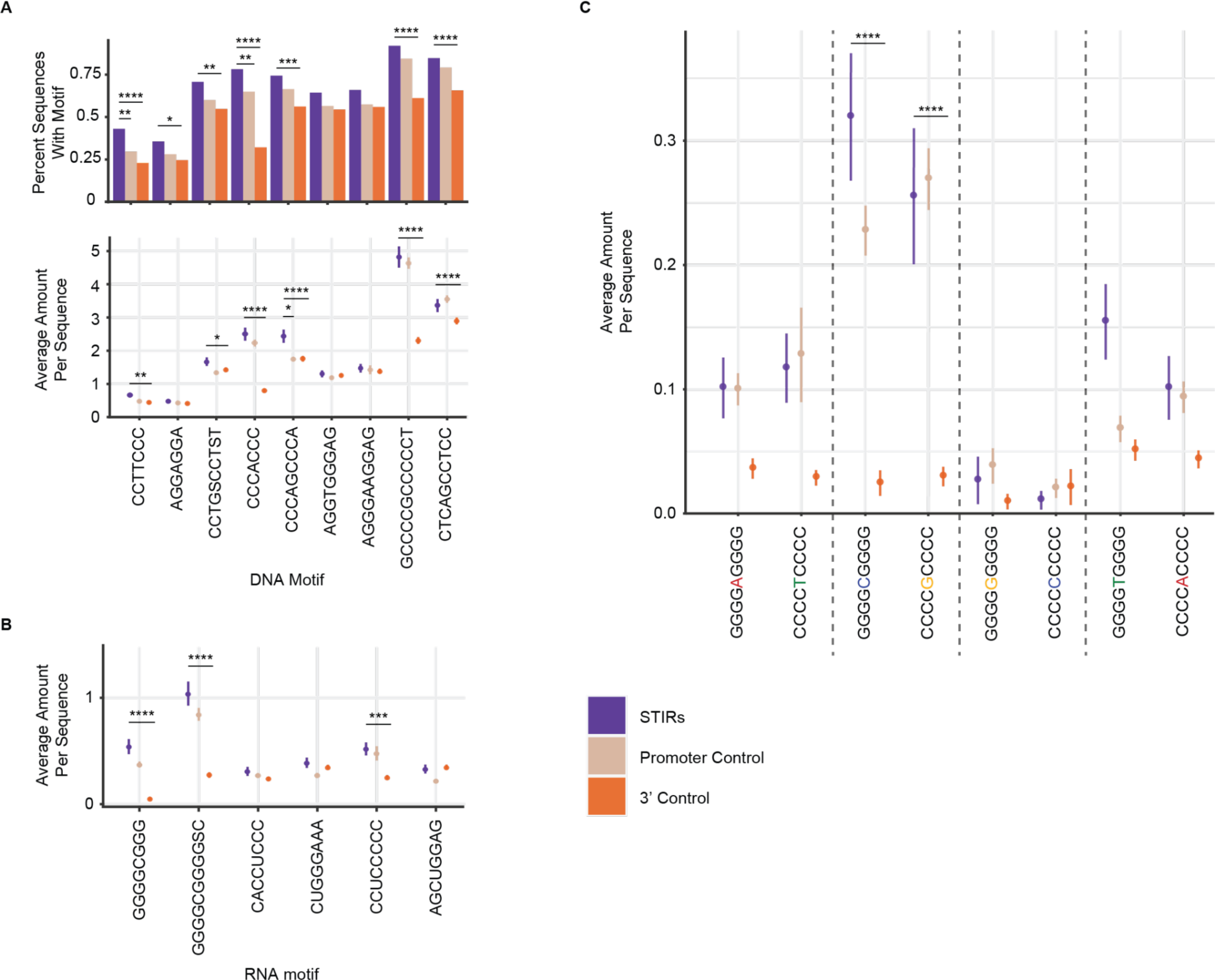
Enriched motifs in STIRs. (A) Top: barplots showing the proportion of STIRs (purple) or control sequences (beige and orange) that carry the DNA-mode STREME-discovered motifs. Shown are Bonferroni corrected proportion test P-values (*:P<=0.05, **:P<=0.01, ***:P<=0.001, ****<=0.0001). Bottom plots show the overall average number and SE of motif occurrences at STIRs or control sequences. P-values were obtained using paired Wilcoxon rank-sum test and corrected using Bonferroni correction. (B) As in the bottom plot of (A), but for the RNA-mode STREME sequence motifs. (C) As in the bottom plot of (A), but for the G4NG4 motifs (or their reverse complement) in HepG2 cells.

**Figure S3.**
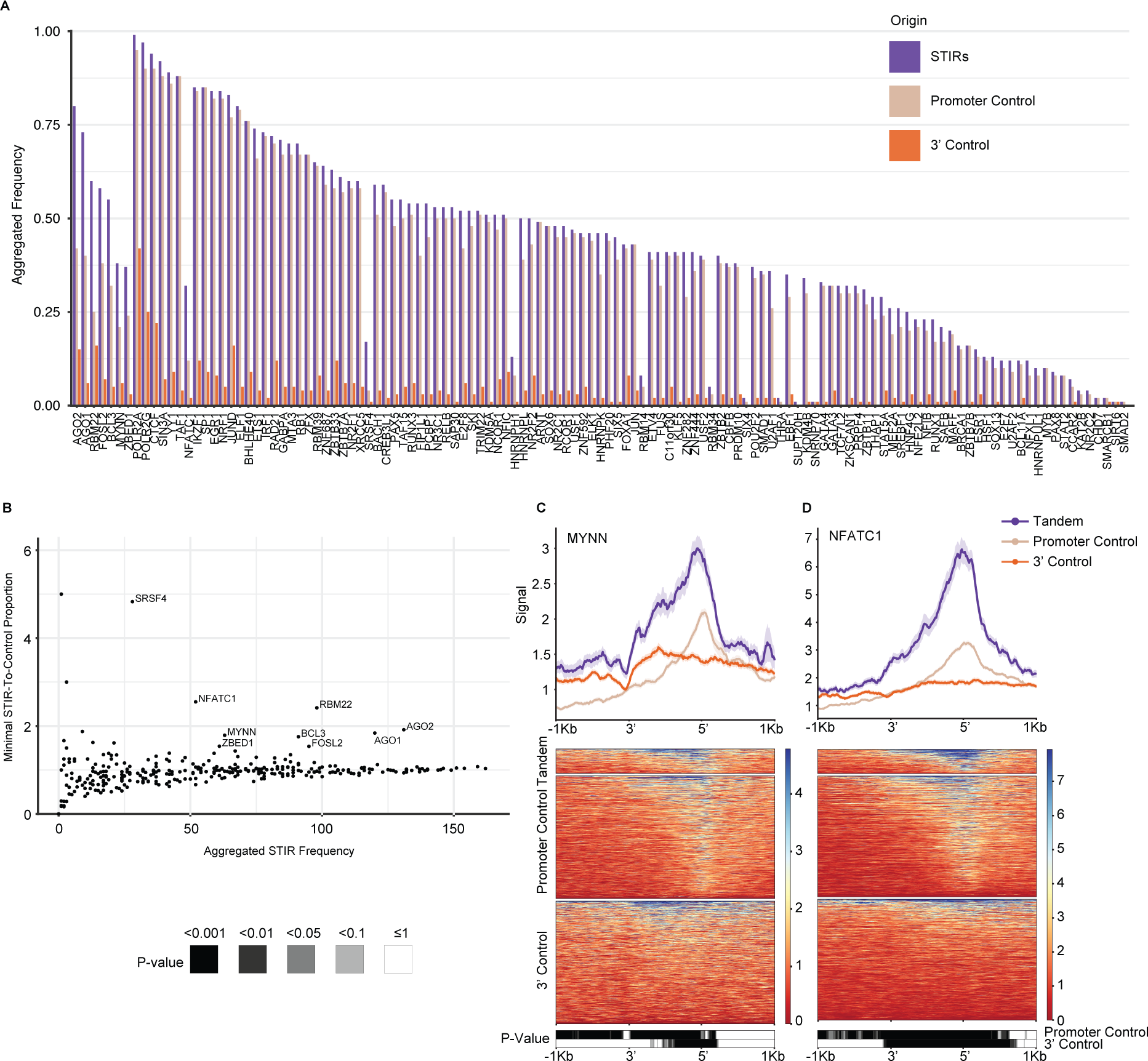
Enriched binding of proteins at STIRs. (A) Barplot showing the proportion of co-expressed STIRs (purple), promoter control- (beige), and 3’ control- (orange) sequences bound by the different proteins (analyzed in K562 cells). Bound sequences are aggregated and counted once in cases multiple binding sites per sequence was suggested. Shown are only proteins with higher binding frequency at STIRs over both controls. Proteins are ordered by STIR ranked frequency and ranked descending order of calculated minimal ratio of frequencies between STIR and each control. (B) Scatter plot showing the number of K562 co-expressed STIRs bound by each protein (as in (A)) versus the minimal tandem-to-control ratio calculated for each control. Indicated are the top enriched proteins at STIRs. (C-D) Metagene analysis (top) and corresponding binding heatmap (center) of median ChIP-seq signal of selected STIR-binding protein candidates MYNN (C), and NFATC1 (D). Bottom heatmap shows the corresponding binned paired Wilcoxon rank-sum tests (Bonferroni corrected).

**Figure S4.**
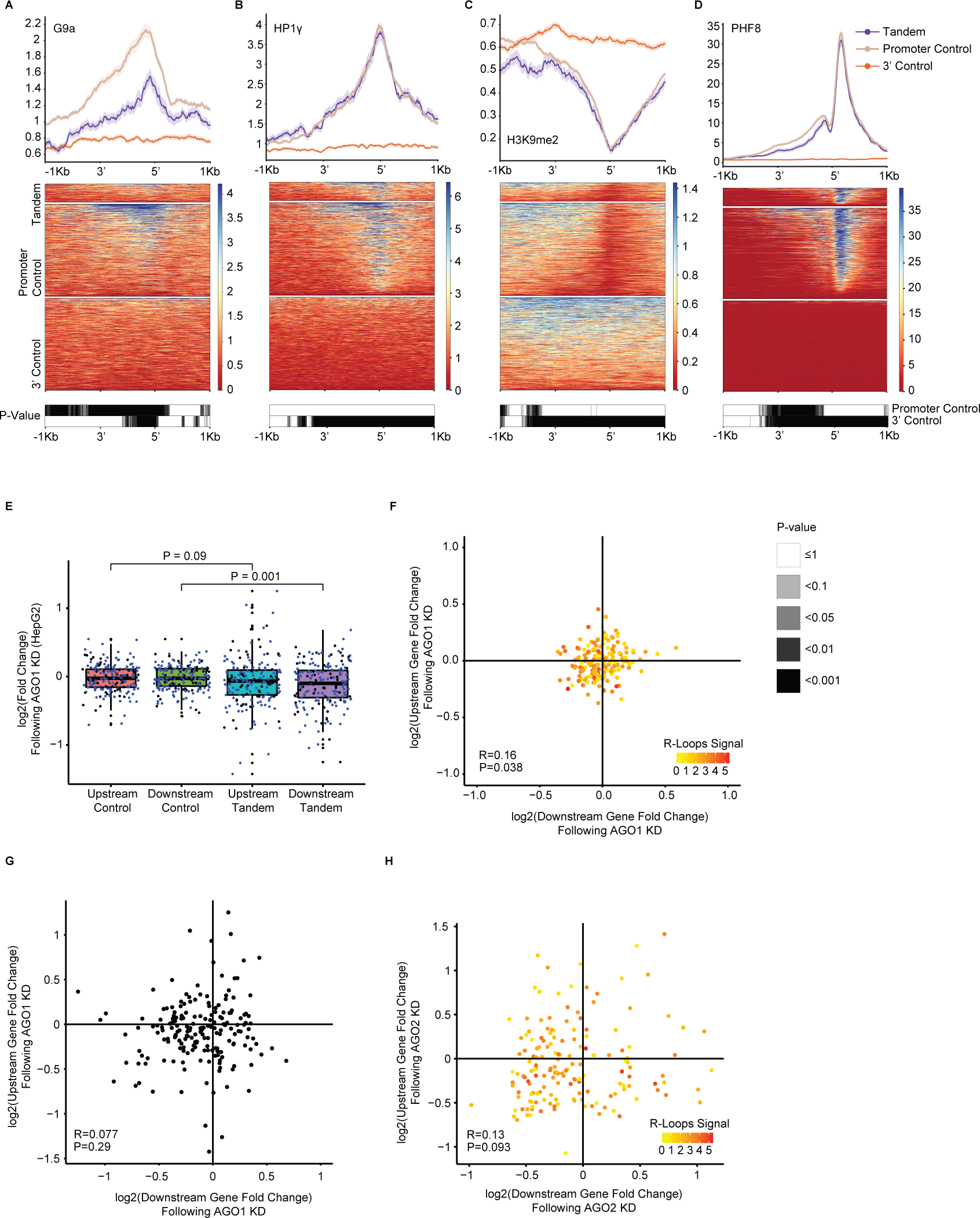
Features promoting the proper expression of tandem gene pairs. (A-D) Metagene analysis (top) and corresponding binding heatmap (center) of median ChIP-seq signal of the H3K9me2-related factors G9a (A), HP1γ (B), H3K9me2 (C) and PHF8 (D). Bottom plots are the corresponding heatmaps of the binned paired Wilcoxon rank-sum tests (Bonferroni corrected). (E) Boxplot of expression changes in upstream or downstream co-expressed tandem genes following AGO1 knockdown (KD) in HepG2 cells (Data from ENCODE). Blue dots correspond to tandem pairs co-expressed in both K562 and HepG2 cell lines (or their respective control). Black dots are tandem genes co-expressed only in the respective cell line. P-values were obtained using paired Wilcoxon rank-sum test. (F) Scatter plot showing the changes in expression following AGO1 KD in K562 cells of each co-expressed tandem pair. Pearson correlation and p-value are indicated. Colors indicate median R-loop signal at the STIR. Spearman correlation between the downstream- or upstream-gene expression changes following KD and median AGO1 R-loop signal was tested, with coefficients of –0.278 and –0.117 and P=3×10^−4^ and P=0.13, respectively. (G) Scatter plot showing the changes in expression following AGO1 KD in HepG2 cells of each co-expressed tandem pair. Pearson correlation and p-value are indicated. (H) Scatter plot showing the changes in expression following AGO2 KD in K562 cells of each co-expressed tandem pair. Pearson correlation and p-value are indicated. Colors represent the median R-loops signal at the intergenic region of each tandem pair plotted. Spearman correlation between the downstream- or upstream-gene expression changes following KD and R-loop median signal at STIRs was calculated and was not significant in both cases (P=0.75 and P=0.37, respectively).

**Figure S5.**
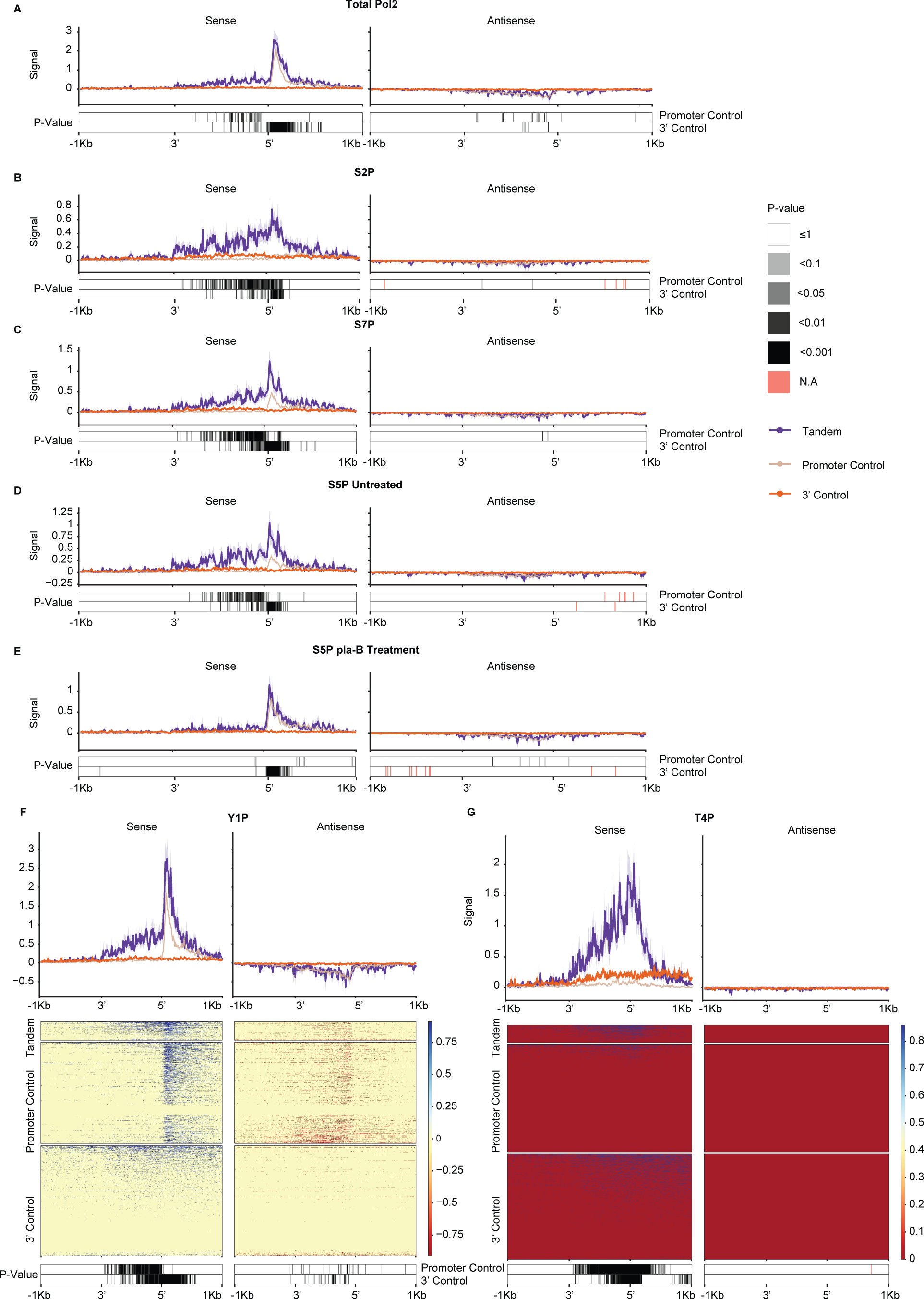
Pol2 marks at STIRs sense and antisense strands. (A-E) Metagene analysis (top) showing the different median Pol2 carboxy-terminal domain modifications (phosphorylated and non-phosphorylated, S2P, S7P, S5P and S5P+Pla-B, respectively) occupancy signal and standard error at the sense and antisense strand of STIRs or controls and their flanking regions. Bottom are the respective corrected paired Wilcoxon rank-sum test P-values. NA P-values are marked in cases where data is missing. Data from Schlackow *et al*., 2017. (F-G) Metagene analysis (top) and corresponding heatmap (center) of median Pol2 CTD modifications (Y1P and T4P, respectively) mNET-seq signal and standard error at the sense and antisense strands of STIRs and controls, and their flanking regions. Bottom are the respective corrected paired Wilcoxon rank-sum test P-values. Data from Schlackow *et al*., 2017.

**Figure S6.**
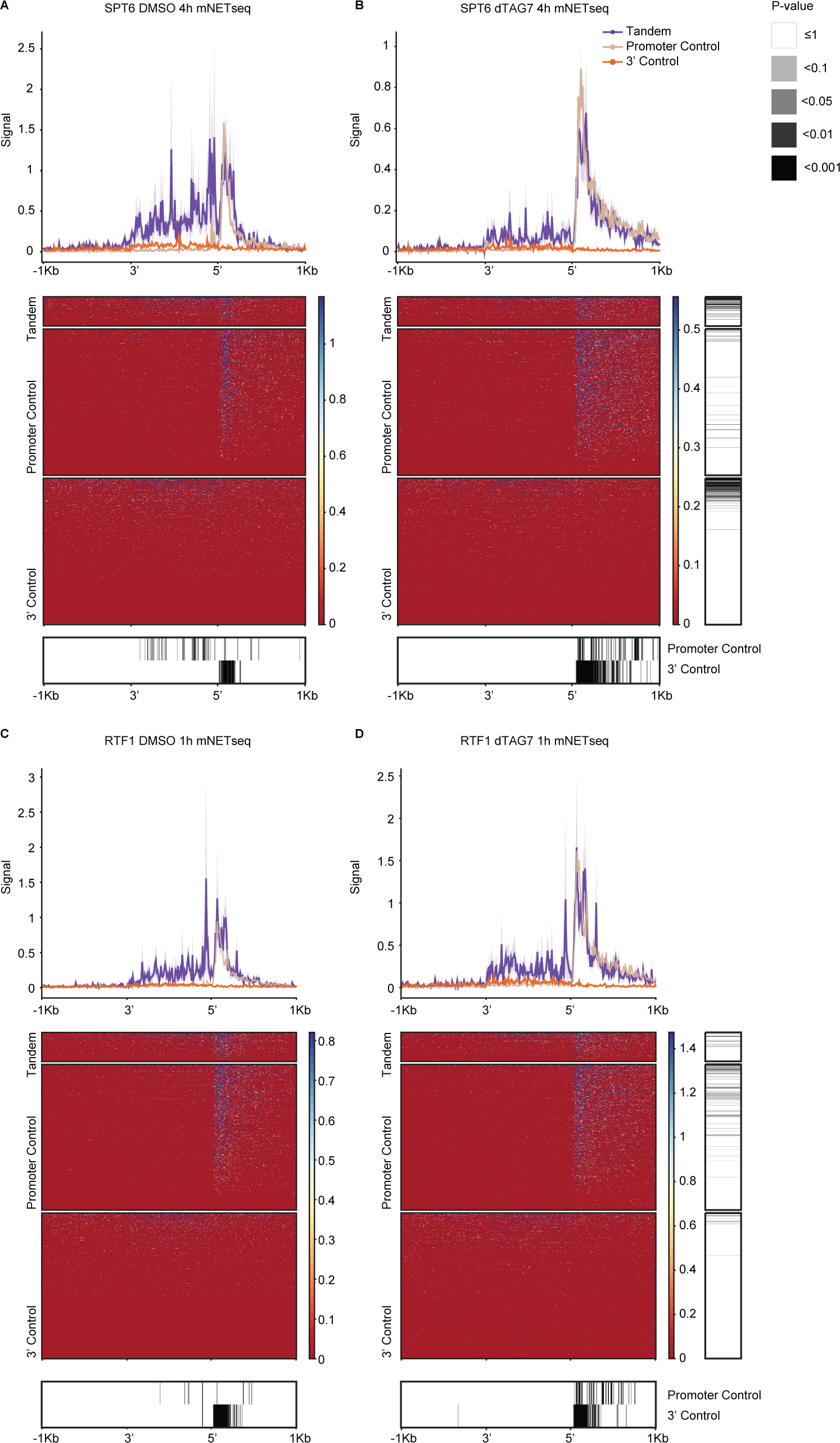
SPT6 degradation is associated with Pol2 paucity at STIRs. (A) Metagene analysis (top) and corresponding heatmap (center) showing mNET-seq median signal following control (DMSO) treatment in K562 co-expressed STIRs and flanking regions and in their controls. Bottom heatmap shows the corrected P-value of binned paired Wilcoxon rank-sum test. Data from Žumer *et al*., 2021. (B) As in (A), for K562 cells 4h after treatment with SPT6 dTAG7. Centered heatmap is ordered by the gene order in the heatmap of (A). Right heatmap is the corrected paired Wilcoxon rank-sum test per gene before and after treatment. Proportion of significantly changed signal for genes following the KD treatment was tested between the group of tandem genes and each control group and was significant in both cases (4.4×10^-31^ and 1.1×10^-9^ for the promoter or the 3’ control, respectively). (C) As in (A), controls for the RTF1 1h treatment with DMSO. (D) As in (C), 1h following RTF1 dTAG7 treatment. Proportion of significantly changed signal for genes following the KD treatment was tested between the group of tandem genes and each control group and had P=0.57 and P=1.6×10^-4^ for the promoter or the 3’ control, respectively.

**Figure S7.**
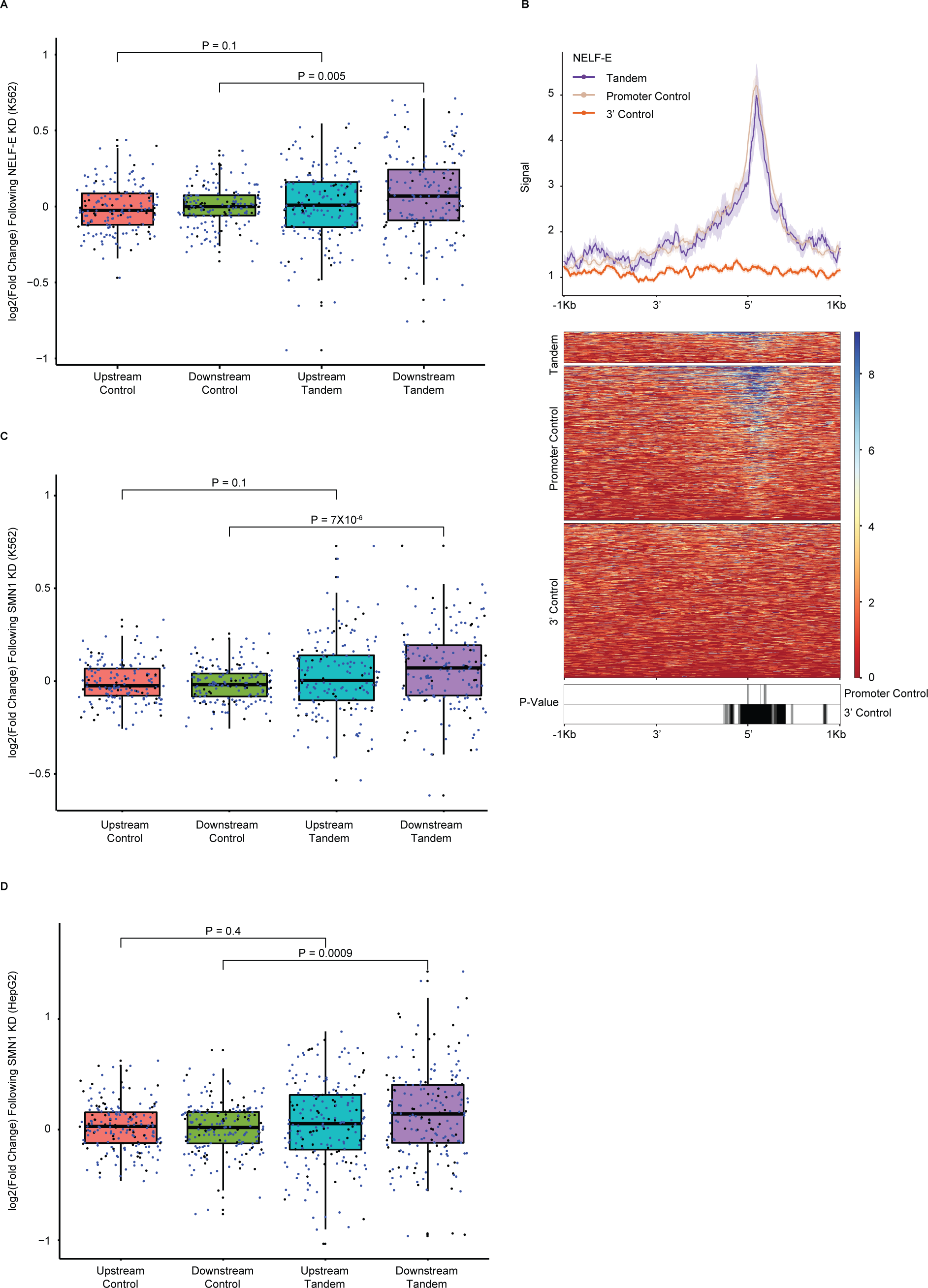
KD candidates for the transcription regulation of downstream tandem genes. (A) Boxplot of expression changes in upstream or downstream co-expressed tandem genes following NELF-E KD in K562 cells (Data from ENCODE) or their controls. Blue dots correspond to tandem pairs co-expressed in both K562 and HepG2 cell lines (or their respective control). Black dots are tandem genes co-expressed only in the respective cell line. P-values were obtained using paired Wilcoxon rank-sum tests. (B) Metagene analysis (top) and corresponding binding heatmap (center) of NELF-E median ENCODE ChIP-seq signal data at K562 co-expressed STIRs and flanking 5’ and 3’ regions and at the control regions. Bottom heatmap shows the binned Bonferroni-corrected paired Wilcoxon rank-sum test P-values. (C) Same as (A), but for SMN1 KD in K562 cell line. (D) Same as (C), but for KD experiment in HepG2 cell line.

**Figure S8.**
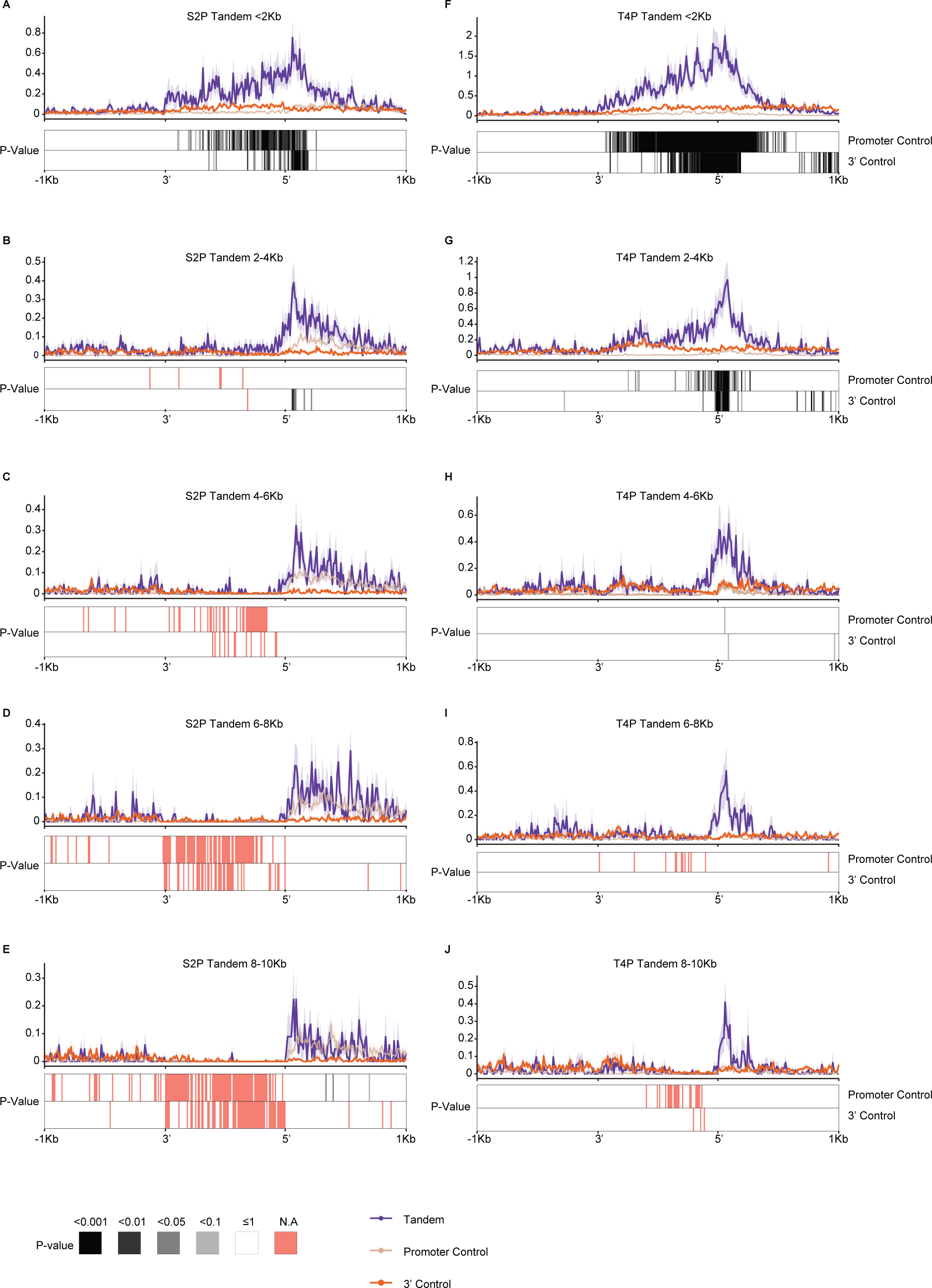
Pol2 marks at increasing lengths of tandem intergenic regions. (A-E) Metagene plot showing Pol2 CTD S2P marks at increasing tandem intergenic lengths (<2kb, 2–4kb, 4–6kb, 6–8kb and 8–10kb) and their respective controls (top). Bottom heatmap shows the binned Bonferroni-corrected paired Wilcoxon rank-sum test P-value heatmap (NA is indicated in cases of missing signal). (F-J) As in (A-E), but for T4P CTD modification.

**Figure S9.**
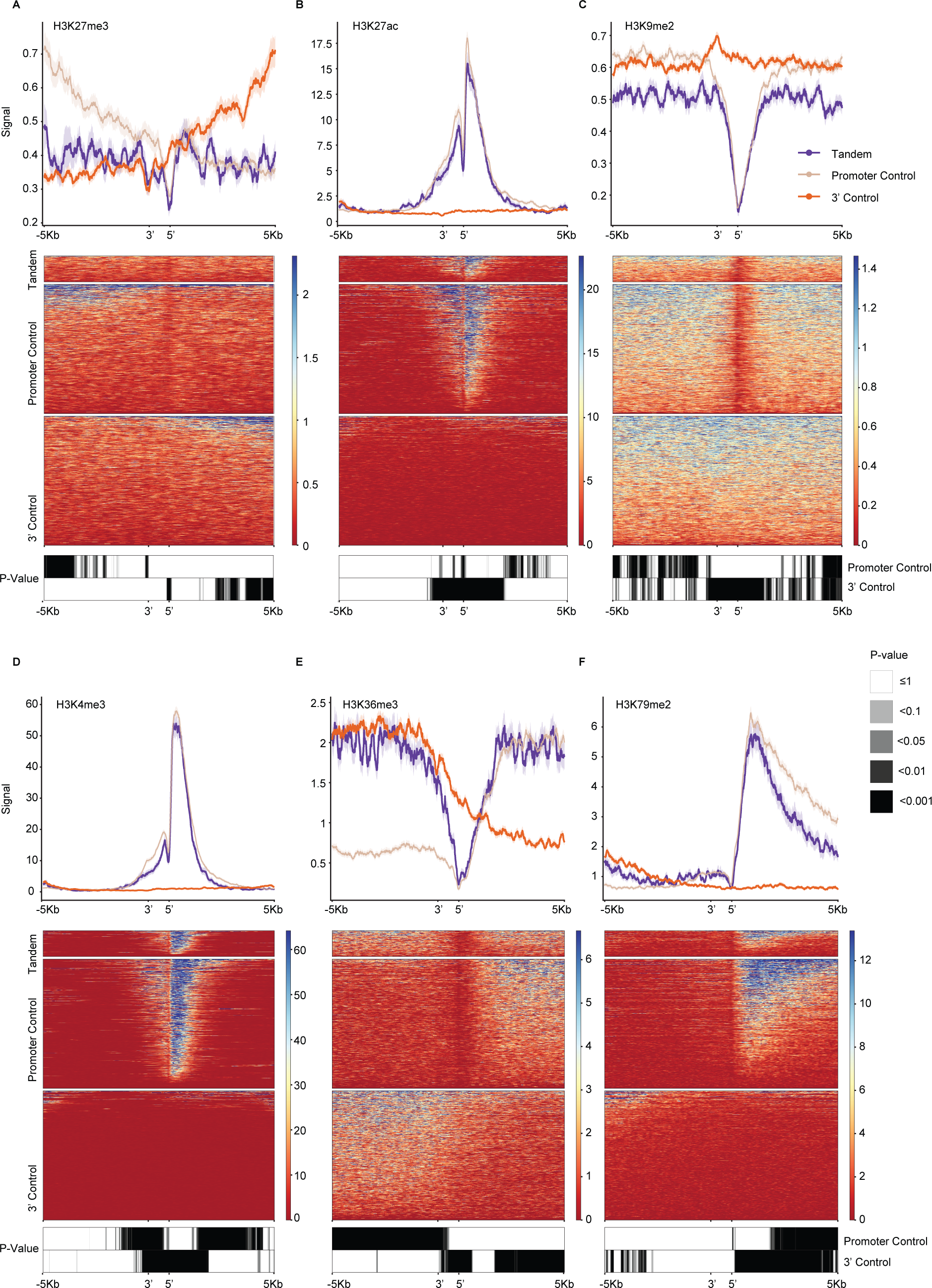
ChIP-seq signal of histone marks at STIRs. (A-F) Metagene plots (top), heatmaps (center) and Bonferroni-corrected binned P-value heatmaps (bottom) of ChIP-Seq signal of different histone modifications H3K27me3 (K562) (A), H3K27ac (K562) (B), H3K9me2 (SK-N-SH) (C), H3K4me3 (K562) (D), H3K36me3 (K562) (E) and H3K79me2 (HepG2) (F). Data from ENCODE.

**Figure S10.**
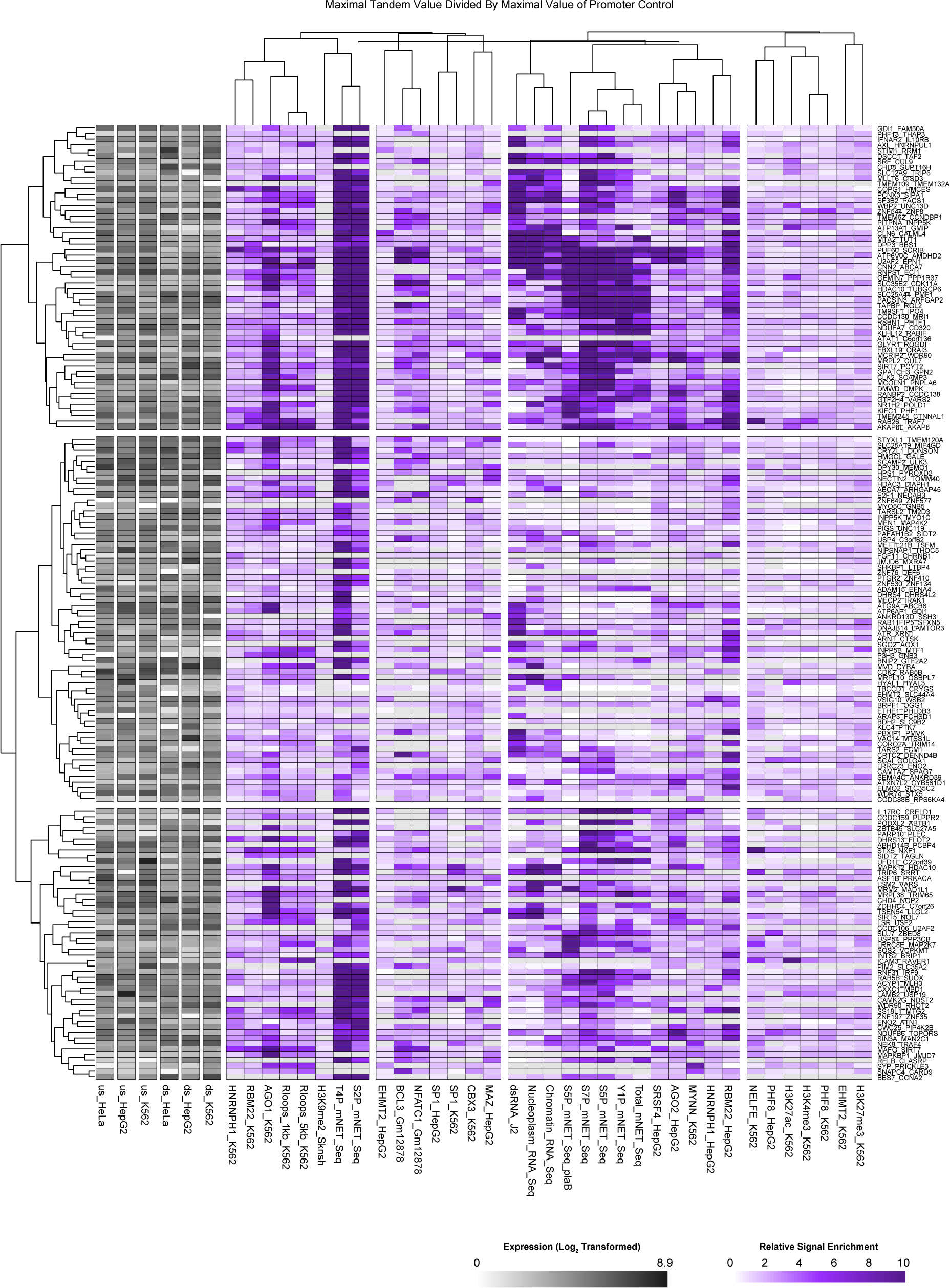
Cluster analysis of STIR normalized to promoter control. Maximal signal per STIR and flanking regions was calculated for tandem genes or their controls (y-axis) per experiment tested in figures 2-8 and S2-S9. Matrix of maximal signal at STIRs was divided by the matching matrix of promoter controls. Clustering of the experiments was done using Pearson’s correlation. Clustering of STIRs was done by computing the Euclidean distance. Left annotations show the log2-transformed expression levels of the respective genes within the different cell lines.

**Figure S11.**
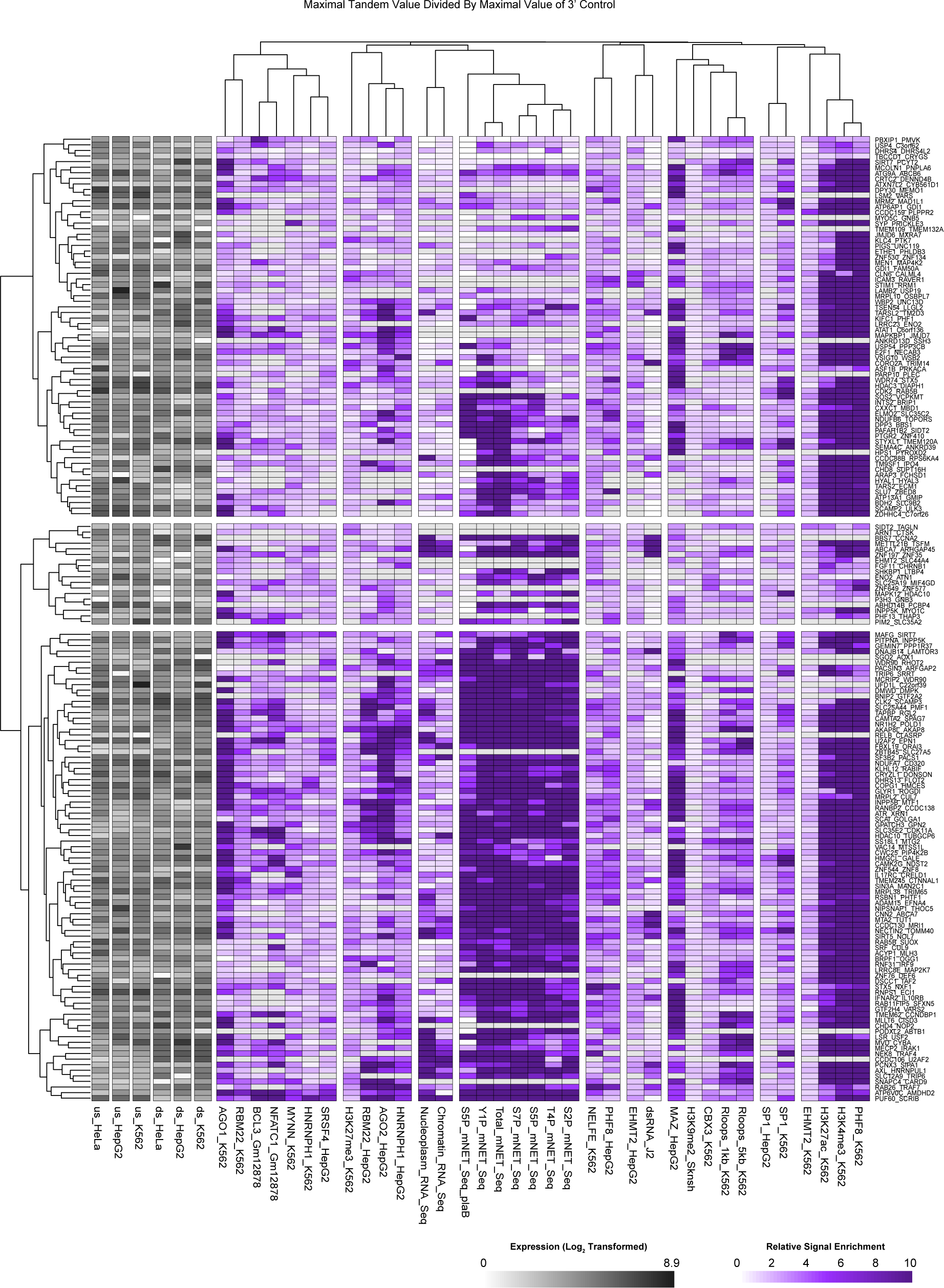
Cluster analysis of STIR normalized to 3’ control. Maximal signal per STIR and flanking regions was calculated for tandem genes or their controls (y-axis) per experiment tested in figures 2-8 and S2-S9. Matrix of maximal signals at STIRs was divided by the matching matrix of 3’ controls. Clustering of the experiments was done using Pearson’s correlation. Clustering of STIRs was done by computing the Euclidean distance. Left annotations show the log2-transformed expression levels of the respective genes within the different cell lines.

